# Positive selection in the genomes of two Papua New Guinean populations at distinct altitude levels

**DOI:** 10.1101/2022.12.15.520226

**Authors:** Mathilde André, Nicolas Brucato, Georgi Hudjasov, Vasili Pankratov, Danat Yermakovich, Rita Kreevan, Jason Kariwiga, John Muke, Anne Boland, Jean-François Deleuze, Vincent Meyer, Nicholas Evans, Murray P. Cox, Matthew Leavesley, Michael Dannemann, Tõnis Org, Mait Metspalu, Mayukh Mondal, François-Xavier Ricaut

## Abstract

Highlanders and lowlanders of Papua New Guinea (PNG) have faced distinct environmental conditions. These environmental differences lead to specific stress on PNG highlanders and lowlanders, such as hypoxia and environment-specific pathogen exposure, respectively. We hypothesise that these constraints induced specific selective pressures that shaped the genomes of both populations. In this study, we explored signatures of selection in newly sequenced whole genomes of 54 PNG highlanders and 74 PNG lowlanders. Based on multiple methods to detect selection, we investigated the 21 and 23 genomic top candidate regions for positive selection in PNG highlanders and PNG lowlanders, respectively. To identify the most likely candidate SNP driving selection in each of these regions, we computationally reconstructed allele frequency trajectories of variants in each of these regions and chose the SNP with the highest likelihood of being under selection with CLUES. We show that regions with signatures of positive selection in PNG highlanders genomes encompass genes associated with the hypoxia-inducible factors pathway, brain development, blood composition, and immunity, while selected genomic regions in PNG lowlanders contain genes related to immunity and blood composition. We found that several candidate driver SNPs are associated with haematological phenotypes in the UK biobank. Moreover, using phenotypes measured from the sequenced Papuans, we found that two candidate SNPs are significantly associated with altered heart rates in PNG highlanders and lowlanders. Furthermore, we found that 16 of the 44 selection candidate regions harboured archaic introgression. In four of these regions, the selection signal might be driven by the introgressed archaic haplotypes, suggesting a significant role of archaic admixture in local adaptation in PNG populations.

## Introduction

After the first arrival of modern humans in New Guinea around 50 thousand years ago (kya) ^1,2^, they rapidly spread across different environmental niches of the island ^3,4^. Since the Holocene (around 11 kya), the Papua New Guinea (PNG) population has been unevenly distributed, with most of the population living at altitude between 1600 and 2400 meters above sea level (a.s.l.) ^5–7^. This population distribution pattern is remarkable considering the challenges PNG highlanders face at this altitude, like the lower oxygen availability to the body ^8^. Studies investigating hypoxic response of the human body in high-altitude populations revealed that selection acted on genes involved in the Hypoxia-Inducible Factor (HIF)-pathway^9,10^, the principal response mechanism to low oxygen at the cellular level. It regulates angiogenesis, erythropoiesis, and glycolysis ^11^. Some high-altitude populations show a limited increase in haemoglobin concentration ^12^ in response to the lower oxygen levels. Indeed, an increase in haemoglobin concentration – as observed in native lowlanders accessing altitude – increases oxygen transport but also results in higher blood viscosity ^13^. In the long term, that process may cause Chronic Mountain Sickness (CMS) and cardiovascular complications ^13^. Interestingly, Tibetan highlanders show selection that is associated with a more restrained increase of haemoglobin concentration at altitude due to increased plasma volume ^14^. This suggests that hypoxia might lead to the selection of a complex haematological response that overcomes the increase in blood viscosity when enhancing oxygen transport. However, the role of selection in response to the environmental challenges by altitude on the genomes of PNG highlanders, who inhabited this environment for the last 20,000 years ^4^, remains mostly unknown. PNG highlanders significantly differ from PNG lowlanders in height, chest depth, haemoglobin concentration, and pulmonary capacities ^15^. Similar differences have been observed between Andean, Tibetan and Ethiopian highlanders and their corresponding lowland populations ^16^. However various factors, like phenotypic plasticity ^17^, diet or physical activities, could explain these phenotype differences. In this paper we explored whether these phenotypes can also be linked to adaptive processes acting on the genome of the PNG highlanders.

Other strong environmental pressures in PNG are infectious diseases (e.g., malaria, dysentery, pneumonia, tuberculosis, etc) that are the leading cause of death in PNG ^18–20^. In this pathogenic environment, malaria stands out among others and could have affected selective pressure in highlanders and lowlanders differently. Incidence of malaria varies enormously between the lowlands and the highlands. While PNG accounted for nearly 86% of the malaria cases in the Western Pacific Region in 2020 ^21^, malaria is practically absent in PNG highlands, possibly because of a limited dispersal of *Anopheles*, the main vector of malaria, at high altitude ^6,22^. It has been suggested that malaria might explain the unbalanced population distribution between PNG highlands and lowlands ^7,23,24^ and thus induces a selection pressure specific to lowlanders. Nonetheless, the period when this specific pathogenic pressure started to impact Papuans remains unclear.

Besides facing these environmental pressures, PNG populations also stand out by their high levels of Denisovan introgression ^25,26^. Denisovan introgressed variant might contribute to Tibetans adaptation to altitude ^27^ and affect the immune system of the PNG population ^28^. Moreover, because some archaic variants show signals of selection among the overall Papuan population ^29–31^, it is conceivable that archaic introgression has contributed to beneficial alleles in PNG populations. However, to date it remains elusive how to which extent archaic introgression contribution to local adaptation varies between PNG populations.

In this study, we identify the genomic regions that show signatures of selection in 54 newly sequenced PNG highlanders and 74 lowlanders. We then screen for the SNP that most likely drives the selection signal in each genomic region under selection. We then explore phenotype associations with candidate SNPs. Finally, we scan selection candidate regions for the presence of introgressed archaic haplotypes and assess the role of introgressed alleles on adaptive processes. Our research provides new insights into local adaptation in PNG populations and its implications on health.

## Material and Methods

### Ethics

This study was approved by the Medical Research Advisory Committee of Papua New Guinea under research ethics clearance MRAC 16.21 and the French Ethics Committees (Committees of Protection of Persons CPP 25/21_3, n_SI: 21.01.21.42754). Permission to conduct research in PNG was granted by the National Research Institute (visa n°99902292358) with full support from the School of Humanities and Social Sciences, University of Papua New Guinea. All samples were collected from healthy unrelated adult donors who provided written informed consent. After a full presentation of the project to a wide audience, a discussion with each individual willing to participate ensured that the project was fully understood.

### Samples

DNA was extracted from saliva samples with the Oragene sampling kit according to the manufacturer’s instructions. Sequencing libraries were prepared using the TruSeq DNA PCR-Free HT kit. About 150-bp paired-end sequencing was performed on the Illumina HiSeq X5 sequencer. We sequenced PNG whole genomes from PNG lowlanders from Daru (n=38, <100 m above sea level (a.s.l)) and PNG highlanders from Mount Wilhelm villages (n=46, 2,300 and 2,700 m a.s.l.) sampled between 2016 and 2019 (EGA accession code XXXXX). To increase our sample size, we included 58 published genomes sampled in Port Moresby, including individuals from different regions in PNG ^3^. We also gained access to PNG whole genome sequences from samples collected at the same sampling places during the same period and sequenced at the National Center of Human Genomics Research (France) or the KCCG Sequencing Laboratory (Garvan Institute of Medical Research, Australia) (unpublished data; F-X. Ricaut personal communication). These additional datasets increased our sample size to a total of 262 PNG whole genomes with 60 individuals from Mount Wilhelm (PNG highlanders), 80 individuals from Daru (PNG lowlanders) and 122 individuals sampled in Port Moresby from different origins (PNG diversity set I) (Note S1, Tables S1-S2). We measured phenotypes associated with body proportion, pulmonary capacities and cardiovascular components in this PNG dataset ^15^ (Note S2, Table S3).

We combined these 262 sequences with published Papuan genomes (n=81, PNG diversity II) ^30,32–35^ and high-coverage genomes from the 1000 Genomes project from Africa (n=207), East Asia (n=202) and Europe (n=190) ^36^ (Note S1).

### Variant Calling

Sequencing data for all samples used in this study were processed together, starting from the raw reads. FASTQ files were trimmed with fastp v0.23.2 ^37^ and converted to BAM using Picard Tools FastqToSam v2.26.2 ^38^. Further processing was performed with Broad Institute’s GATK Germline short variant discovery (SNPs and Indels) Best Practices ^39^. HaplotypeCaller tool was used to produce individual sample GVCF files, which were further combined by JointGenotyping workflow to create multi-sample VCF files. GATK v4.2.0.0 was used ^40^. Data were processed with GRCh38 genome reference (Note S3).

### Filtering

Unless otherwise stated, we performed the analysis on biallelic SNPs with a maximal missing rate of 5% that remained after genomic masking (Note S7). For each pair of related individuals to the second degree, when relevant, we kept the individuals with the highest number of phenotypes measurements or the individual with the highest mean of coverage. We removed two PNG samples with low call rate from any further analysis. Quality and kinship filtering resulted in 249 unrelated genomes among the PNG highlanders, lowlanders and the PNG diversity set I: 54 sequences of PNG highlanders, 74 sequences from PNG lowlanders and 121 sequences from individuals originating from different parts of PNG and sampled in Port Moresby (PNG diversity set I; Notes S1, S4-S7, Tables S1-S4, Figures S1-S2). The unrelated and filtered dataset also includes 262 published Papuan sequences (n=81, PNG diversity II) ^30,32–35^ and sequences from the 1000 Genomes project from Africa (n=207), East Asia (n=202) and Europe (n=190) ^36^ (Note S1).

### Population structure

Principal Component Analysis (PCA) was performed on the unrelated dataset filter for variant with minor allele frequency <5% and pruned for linkage disequilibrium (Note S8) using the smartpca program from the EIGENSOFT v.7.2.0 package ^41^. To prune variants in high linkage disequilibrium, we used PLINK v.1.9 using the default parameters of 50 variants count window shifting from five variants and a variance inflation factor (VIF) threshold of 2 ^42^. The LD pruned dataset included 469,584 SNPs (4,809,440 SNPs before pruning).

We used the R-3.3.0 software to plot the PCA. We computed the PCA to the tenth principal component. We ran ADMIXTURE v1.3 ^43^ on the same dataset from components K=2 to K=6. To define how many components composed the most likely model, we computed each component’s confidence interval of the cross-validation error by repeating it 50 times (Note S9).

### Phasing

We phased genomes from Mt Wilhelm, Daru, PNG diversity set I, Africa, Asia and Europe using shapeit4 (v4.2.2) ^44^. We phased the samples statistically without reference, as the reference haplotypes panel for the PNG population does not exist (Note S10).

### Selection analysis

We aimed to identify genomic regions carrying signatures of positive selection in PNG highlanders and lowlanders using three metrics. We computed Population Branch Statistic (PBS), a method based on allele frequency, to detect recent natural selection signals in PNG highlanders and lowlanders ^45^ (Note S11). For the PBS scores in PNG highlanders, we used PNG lowlanders as reference and Yorubas (YRI) from 1000 Genome as the outgroup. When performing PBS on PNG lowlanders, we used PNG highlanders as reference and the YRI as the outgroup. In both cases, we obtained a PBS score for every biallelic SNP. We then defined sliding windows of 20 SNPs with a step of 5 SNPs to identify multiple adjacent SNPs with an elevated PBS score (which lowers the random chances due to drift). We assigned the average PBS score of all the SNPs included in the sliding window as the PBS score of the window. We kept the sliding windows with an average PBS score in the 99^th^ percentile and merged the top sliding windows that are 10kb maximum from each other. The top PBS score of the sliding windows in the region was given to the whole merged region.

In addition, we computed the cross-extended haplotype homozygosity (XP-EHH) ^46^ on the phased dataset with selscan (v2.0.0) ^47^ to test for positive selection using haplotype information (Note S12). We computed XP-EHH using PNG highlanders as the target population and PNG lowlanders as the reference population. While the maximal scores define regions under selection in PNG highlanders, the lowest scores indicate the regions under selection in PNG lowlanders. We determined the top SNPs for XP-EHH score in PNG highlanders as the SNP with XP-EHH score in the 99^th^ percentile. We kept the SNPs with XP-EHH score in the 1st percentile for PNG lowlanders. We merged these top SNPs in windows: two top SNPs distant by at most 10kb are included in the same window. This merging step results in windows whose endpoints are the two most distant top SNPs included in the window.

Next, we combined the PBS and XP-EHH scores in a Fisher score ^48^ (Note S13). We used the sliding windows of 20 SNPs, and 5 SNPs step defined for the PBS score. For each of these sliding windows, we gave as XP-EHH score the highest XP-EHH score among the 20 SNPs included in the windows. We combined the PBS and XP-EHH scores in a Fisher Score (-*log*_10_(*PBS_percentilrank_*) - *log*_10_(*XP - EHH_percentilrank_*) ^48^ for each sliding window. Finally, we selected the windows Fisher Score in the 99^th^ percentile and merged them when they were distant of maximum 10kb. We extended the top 10 merged windows with the highest score for each of the three methods by a 50kb flanking region. Finally, we merged the overlapping regions from these 30 top regions to obtain the final non-overlapping regions of interest that we will use further.

Because of the low number of individuals per population in the PNG diversity sets I and II and the high genetic diversity in PNG (Figures S3-S4), we did not include these samples in the selection analyses described above.

### Selection of the SNPs of interest

We computed ancestral recombination graphs for the phased dataset with Relate (v1.1.8) ^49^ (Note S14). We generated coalescence rates through time within PNG highlanders and lowlanders from their respective subtrees. Finally, we extracted the local tree for each SNP in the regions of interest from PNG highlanders and lowlander subtrees. We used these local trees as input for Coalescent Likelihood Under Effects of Selection (CLUES) (v1) ^50^ (Note S15). CLUES assigns a likelihood ratio (logLR) to each SNP of interest that reflects the support for the non-neutral model. For each SNP in the region of interest, we computed logLR five times by re-sampling the local tree branch length and averaged the logLR for the five runs. To decide between the top five SNPs with the higher average logLR in each genomic region, we generated the logLR 50 additional times for these five SNPs. We considered the SNP with the highest average log LR after 50 runs as the SNP the most likely to drive selection within the regions under selection (aka candidate SNPs). Because SNPs with low DAF (Derived Allele Frequency) are unlikely to be under selection, we did not consider SNPs with DAF lower than 5%. We also filtered out fixed variants for which CLUES cannot compute the logLR.

### Association in the UK biobank

To further understand how the candidate SNPs affect phenotypes, we downloaded the UK biobank’s summary statistics ^51^ for the 1,931 phenotypes with more than 10,000 samples (Note S17). We extracted the p-value and the beta of the candidate SNPs for each phenotype. To avoid the ancestry sample size bias present in UKBB, we only extracted the p-value (pval_EUR) and beta score (beta_EUR) for European ancestry. Because the PNG population has a unique genetic diversity absent in Europeans, some candidate SNPs were not listed in the UK biobank. In that case, we looked for summary statistics for the closet SNP from a 1kb upstream and 1kb downstream region. After extracting the SNP summary statistics for every phenotype, we only consider the phenotype of interest if the log(p-value) is lower than −11.29 to correct for multiple testing considering the significance threshold of log(10^-8^) that needs to be corrected for the number of phenotypes studied 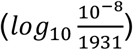. Finally, we corrected the orientation of the beta value from the alternative allele to the derived allele.

### Association test

We used Genome-wide Efficient Mixed Model Association (GEMMA) (v0.98.4) ^52^ to detect if the candidate SNPs are associated with any phenotypes that we measured in the PNG highlanders, lowlanders and PNG diversity set I datasets (Note S16). As we did previously ^15^, we corrected the haemoglobin concentration, blood pressure, heart rate and BMI for age and gender and the chest depth, waist circumference, weight, and pulmonary function measurements (FEV1, PEF and FVC) for age, gender and height using a multiple linear regression approach.

We performed association tests with a univariate Linear Mixed Model (LMM) for the SNPs of interest and each corrected phenotype. To increase our sampling size, we performed these association tests using all the PNG individuals (highlanders, lowlanders and PNG diversity set I) with at least one phenotype measurement (n=234) (Table S3). We incorporated into the LMM the centred relatedness matrix computed with GEMMA using all the 234 PNG sequences to correct for population stratification. We corrected each p-value for the number of SNPs tested with the Benjamini-Hochberg procedure ^53,54^. Because these phenotypes can be gathered in five groups of highly correlated phenotypes ^15^, we used a threshold for significance of 0.01 (0.05/5) to correct for the number of phenotypes tested.

### Introgression

To reveal similarities between PNG haplotypes and archaic haplotypes for the genomic regions under selection in PNG highlanders and lowlanders, we used haplostrips (v1.3) ^55^ within PNG, African, Asian and European samples with Altai ^56^ Neanderthal or Denisovan ^57^ genome as reference haplotypes (Note S18). We explored archaic allele frequencies in the Papuans from the SGDP dataset ^34^ in the regions with introgressed haplotypes in PNG highlanders and lowlanders. We calculated these frequencies on aSNPs, which were defined to be SNPs with one allele (i) present in at least PNG high- or lowlander, (ii) found in a homozygous state in one of the three archaics of the Altai, Vindija Neanderthals and Denisovan ^56–58^ and (iii) being absent in the 1,000 Genomes YRI population.

### Prediction of variant effect

As an additional effort to decipher the function of the candidate SNPs (e.g. gene expression or changes in protein sequence), we looked for significant eQTLs for each candidate SNP using the Genotype-Tissue Expression (GTEx) Portal ^59^. In addition, we downloaded the 111 reference human epigenomes from the Roadmap epigenomics project ^60^ to explore which chromatin state the candidate SNPs fall in different tissue types. Finally, we used The Ensembl Variant Effect Predictor (VEP) ^61^ on the region under selection to detect missense variants in these regions with the canonical flag.

## Results and discussion

### Selection scans results in PNG highlanders and PNG lowlanders

To study selection specific to PNG highlanders or PNG lowlanders, we used 54 newly sequenced genomes from three villages in PNG Highlands located in Mount Wilhelm between 2,300 and 2,700 meters above sea level (a.s.l.) and 74 newly sequenced genomes from Daru island (<100 m a.s.l.). We computed frequency-based (PBS) and haplotype-based (XP-EHH) selection statistics – two selection tests based on distinct genetic signatures – to detect candidate regions for selection in PNG highlanders and lowlanders. Both selection statistics require a target and reference population, allowing us to identify the signal of selection within the target population (PNG highlanders or PNG lowlanders) but absent in the reference population (PNG lowlanders or PNG highlanders, respectively). We also combined both these statistics in a Fisher Score ^48^ to detect the region with extended haplotype homozygosity and carrying multiple variants with high allele frequency. For each selection statistic (PBS, XP-EHH and Fisher Score), we kept the ten regions with the highest score leading to 30 genomic regions of interest for PNG highlanders and lowlanders (Tables S5-S6). We merged the overlapping regions between methods, resulting in a final number of 21 regions of interest in PNG highlanders (Tables 1, S5, Figure 1) and 23 in PNG lowlanders (Tables 2, S6, Figure 1).

**Figure 1:**
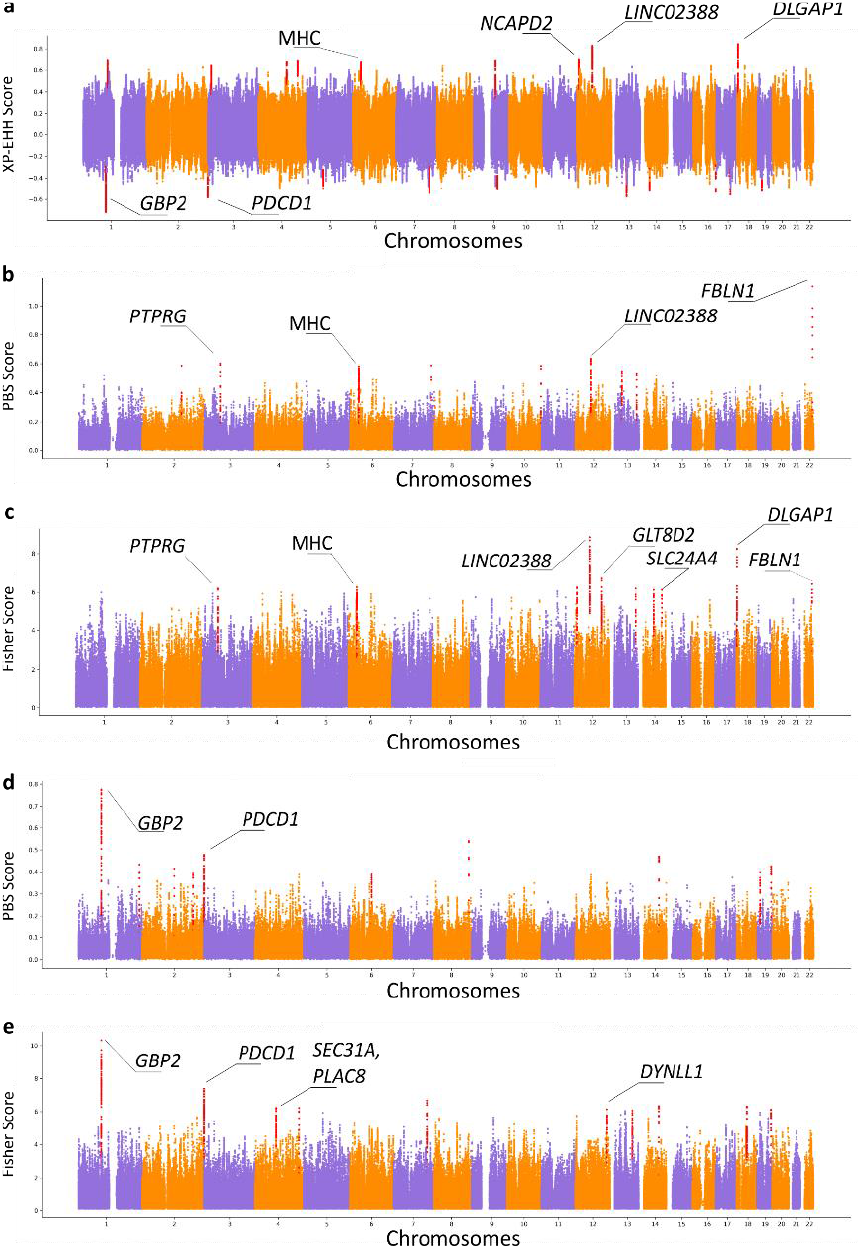
Manhattan plots for the three selection scans among PNG highlanders and lowlanders. Candidate genes discussed in the paper are shown. **(a)** XP-EHH scores using PNG highlanders as the target population and PNG lowlanders as the reference population. Genomic regions with the highest score indicate selection in PNG highlanders. Genomic regions with the lowest score indicate selection in PNG lowlanders. **(b)** PBS scores using PNG highlanders as the target population, PNG lowlanders as the reference population, and Yorubas from 1000G as the outgroup. **(c)** Fisher Scores combining the PBS and XP-EHH scores of PNG highlanders. **(d)** PBS scores using PNG lowlanders as the target population, PNG highlanders as the reference population, and Yorubas from 1000G as the outgroup. **(e)** Fisher Scores combining the PBS and XP-EHH scores of PNG lowlanders.

**Table 2:** Merged regions under selection and SNP most likely to be selected in PNG highlanders.

| Merged top regions | Score | Protein coding genes in the region | Archaic introgression | Candidate SNP for the region | DAF | Significant association (UK Biobank) | Distance to the closest SNP in UKBB (BP) |
| --- | --- | --- | --- | --- | --- | --- | --- |
| chr1:95529290-95736826 | XPEHH | . | Denisova | rs887476833-G/A | 0.55 | -* | + 33 |
| chr2:151012094-151201575 | PBS | . | . | rs74621527-G/A | 0.92 | - | 0 |
| chr3:13010340-13217789 | XPEHH | **IQSEC1** | Denisova | rs374181005-T/C | 0.41 | -* | - 50 |
| chr3:61779523-62009858 | PBS, Fisher | **PTPRG** | . | rs79600167-G/A | 0.77 | - | 0 |
| chr4:110182324-110384099 | XPEHH | **ELOVL6** | Altai Neanderthal | rs943845085-G/A | 0.42 | -* | - 22 |
| chr4:152704503-152970509 | XPEHH | *TIGD4, ARFIP1, FHDC1* | . | rs369030953-A/G | 0.59 | - | - 88 |
| chr6:30916070-31153184† | XPEHH | *VARS2, SFTA2, MUCL3, MUC21, MUC22, HCG22, C6orf5, PSORS1C1, CDSN, PSORS1C2, PSORS1C1, CCHCR1* | . | rs940110341-G/A | 0.61 | Blood composition* (Table S12) | + 94 |
| chr6:33006055-33132312† | PBS, Fisher | *HLA-DAO, HLA-DPA1, HLA-DPB2* | . | rs9277772-T/C | 0.21 | Body proportion, blood composition, other phenotypes (Table S11) | 0 |
| chr7:147590904-147718219 | PBS | **CNTNAP2** | . | rs17170618-T/C | 0.52 | - | 0 |
| chr9:85458922-85745092 | XPEHH | **AGTPBP1** | . | rs28728004-C/A | 0.69 | Other phenotypes* (Table S12) | - 7 |
| chr10:131112245-131235951 | PBS | **TCERG1L** | Altai Neanderthal | rs10829909-T/G | 0.43 | - | 0 |
| chr12:6452552-6662260 | XPEHH | *TAPBPL, VAMP1, MRPL51, GAPDH, NOP2, LPAR5, ING4, ACRBP, CHD4, IFFO1, NCAPD2* | Denisova | rs74576183-A/G | 0.71 | Blood composition (Table S11) | 0 |
| chr12:9886812-10055333 | Fisher | *KLRF2, CLEC2A, CLEC12A, CLEC1B, CLEC12B, CLEC9A* | . | rs536947-C/T | 0.91 | - | 0 |
| chr12:58391529-58634980 | XPEHH, PBS, Fis | . | Altai Neanderthal | rs376870800-A/G | 0.70 | -* | - 160 |
| chr12:103783315-104121479 | Fisher | **NT5DC3, HSP90B1, GLT8D2, HCFC2, NFYB, TDG** | Denisova | rs1032698711-G/A | 0.47 | -* | - 22 |
| chr13:47639988-47825193 | PBS | . | . | rs1033760372-C/A | 0.19 | -* | - 34 |
| chr13:104734734-104875020 | PBS, Fisher | . | Denisova | rs16965509-G/A | 0.50 | - | 0 |
| chr14:60157772-60377317 | Fisher | **PCNX4, DHRS7, PPM1A** | Denisova | rs1033848215-A/G | 0.32 | Other phenotypes* (Table S12) | - 2 |
| chr14:92230479-92401520 | Fisher | **SLC24A4** | . | rs8003454-C/T | 0.52 | - | 0 |
| chr18:4072997-4251153 | XPEHH, Fisher | **DLGAP1** | Altai Neanderthal | rs371858795-G/A | 0.77 | Other phenotypes* (Table S12) | + 124 |
| chr22:45519818-45644906 | PBS, Fisher | **FBLN1** | . | rs1601558750-G/A | 0.10 | Body proportion* (Table S12) | + 101 |

Genomic coordinates are given for GRCh38 DAF is given for PNG lowlanders. †Reference Assembly Alternate Haplotype Sequence Alignments \*SNP not present in the UK Biobank, we look association for the closest SNP within 1KB upstream and downstream region Genes in bold are the closest to the candidate SNP defined with CLUES for the region Putative introgressed regions are given using haplostrips

**Table 2:**
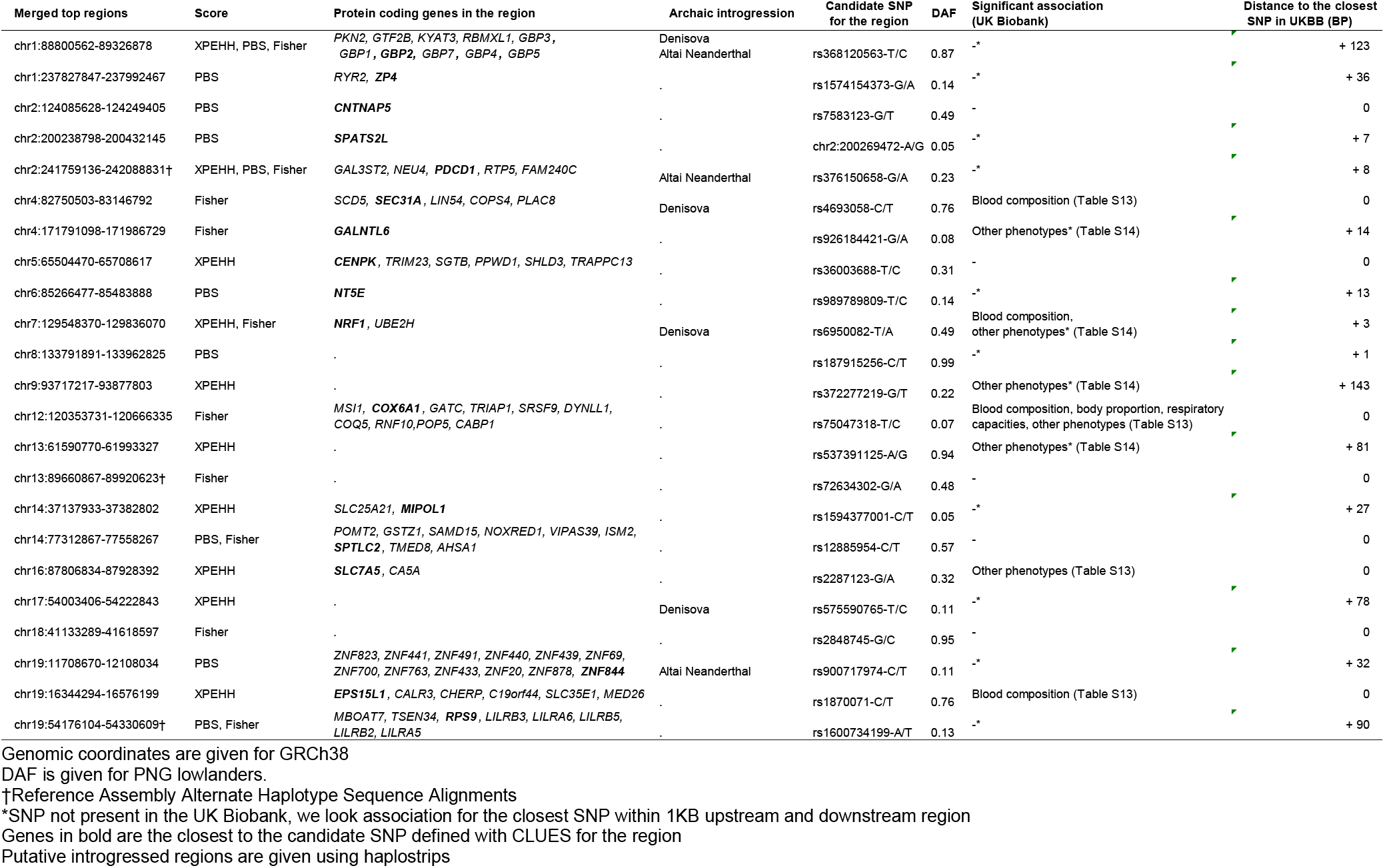
Merged regions under selection and SNP most likely to be selected in PNG lowlanders.

The 21 regions showing signatures of selection in PNG highlanders encompass 54 genes, including genes involved in the regulation of platelet adhesion (ex: *FBLN1* ^62^), HIF-pathway (ex: *LINC02388* ^63^), neurodevelopment (ex: *DLGAP1* ^64^) and immunity (ex: MHC locus ^65^) (Tables 1, S5, Figure 1). The region with the highest Fisher score and second highest PBS and XP-EHH scores in PNG highlanders includes the long intergenic non-protein coding RNA *LINC02388*. This intergenic RNA is associated with the serum levels of protein LRIG3 ^63^ that impact angiogenesis – the formation of new blood vessels – in glioma cells through regulation of the HIF-1α/VEGF pathway ^66,67^. Comparably to other axes of the HIF pathway under selection in high-altitude populations ^9,10^, we hypothesise that this selection signature on *LINC02388* might reflect adaptive processes counteracting hypoxia by affecting the formation of new blood vessels. This axis of the HIF pathway might maintain oxygen transport to appropriate levels in PNG highlanders while limiting the increase in haemoglobin concentration and blood viscosity. Moreover, five of the ten regions with the highest Fisher score include a gene associated with cardiovascular phenotypes *(FBLN1* ^62^*, GLT8D2* ^68^, *DLGAP1* ^69^, *PTPRG* ^70^ and *SLC24A4* ^71^). This observation supports our hypothesis that selection in PNG highlanders acted on genes that might have helped them to counteract the hypoxic condition of their environment.

Genomic selection candidate regions in PNG lowlanders encompassed multiple immunity-related genes *(PLAC8* ^72^, *SEC31A* ^73^, *PDCD1* ^74^, *DYNLL1* ^75^) (Tables 2, S6, Figure 1). Notably, the region with the highest XP-EHH, PBS and Fisher Score includes several genes from the guanine-binding protein family (GBP). This gene family is associated with protective effects against diverse pathogens ^76^. The lowlander-specific selection signature for this gene family, supports the hypothesis that adaptive processes in this population were linked to the specific pathogenic pressure PNG lowlanders faced.

### Selected SNPs phenotypic associations

Next, we sought to identify the most likely selection target SNPs in each candidate region. To this end we reconstructed allele frequency trajectories through time for all the SNPs in a candidate region for selection for the last 980 generations (27,440 years), using CLUES ^50^ and selected the SNP with the largest average log(LR) (here onwards they will be regarded as candidate SNPs; Tables 1-2, S7-S10). Next, we applied two complementary approaches to explore the phenotypic effects of each candidate SNPs. First, we queried GWAS summary statistics from the UK Biobank for each candidate SNP. Seven candidate SNPs of PNG highlanders (or the closest SNPs when the candidate SNP was not present in the UK Biobank) demonstrate significant association with at least one phenotype of the UK Biobank (Table 1, Table S11-S12). Three of these SNPs are significantly associated with haematological phenotypes. Similarly, among PNG lowlanders, eight candidate SNPs show significant associations in the UK Biobank and four with haematological phenotypes (Table 2, Table S13-S14).

We were able to replicate associations of these SNPs under selection and cardiovascular components using phenotypes measurement done for PNG highlanders, lowlanders and PNG diversity set I datasets. After correction for age, gender and the number of tested SNPs, we identified two significantly associated SNPs, both of which showed associations with heart rate (pval_adjusted_ < 0.05; pval adjusted for the number of SNPs tetsed) (Figure 2) although this association does not survive after correcting the significance threshold for the number of tested phenotypes (pval_adjusted_ > 0.01) (Note S16, Table S15). The derived allele G of rs74576183-A/G, an intronic variant of *NCAPD2*, that is under positive selection in PNG highlanders based on CLUES results (Table S7) might be associated with a slower heart rate (pval_adjusted_= 0.046, beta=-2.981; Table S15, Figure 2). On the contrary, the derived allele T of rs4693058-C/T, an intronic variant of *SEC31A*, that is under positive selection in PNG lowlanders (Table S8) might be associated with a faster heart rate (pval_adjusted_= 0.046, beta=3.137; Table S15, Figure 2). Interestingly, these two SNPs showed significant associations with diverse haematological phenotypes in the UK biobank as well (Tables S11, S13). It is possible that these associations with heart rate might reflect an association with other haematological components that were not measured in the PNG samples. Indeed, heart rate correlates with haematological components that are usually overlooked and might be the real target of selection ^14^.

**Figure 2:**
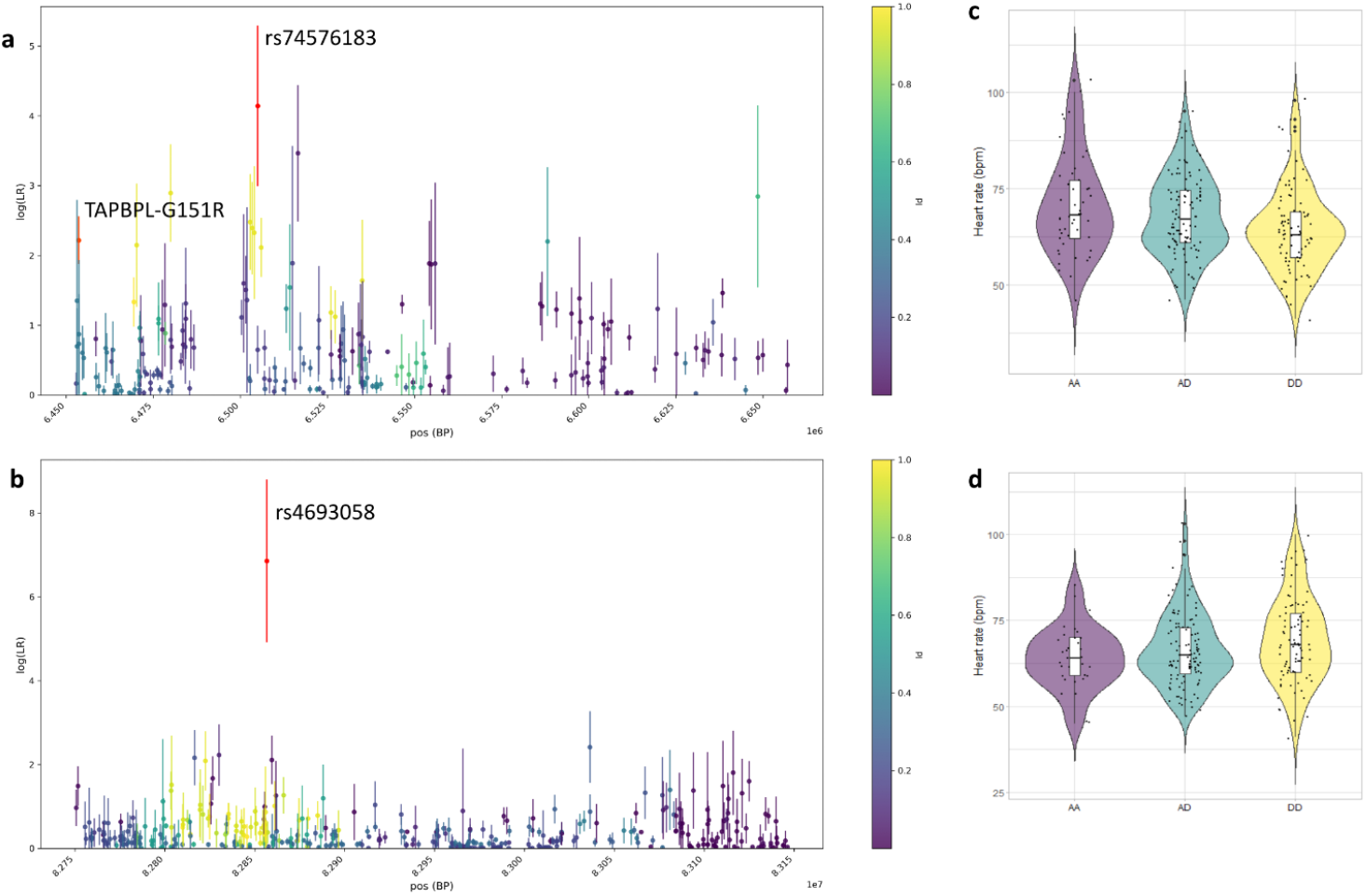
**a, b log(LR) for SNPs in regions under selection** after 5 runs of CLUES or 50 runs of CLUES for each of the five top SNPs for the candidate region. Candidate SNP driving selection for the region are shown in red. Colour scale indicates linkage disequilibrium with the candidate SNP. (**a**) Region chr12:6452552-6662260, that is under selection in PNG highlanders. Candidate SNP for the region is rs74576183-A/G. Missense variant (TAPBPL-G151V) in high LD with rs74576183-A/G is shown in orange. (**b**) Region chr4:82750503-83146792, that is under selection in PNG lowlanders. Candidate SNP is rs4693058-C/T. **c, d Violin plot of the heart rate distribution in PNG depending of their genotype for the candidate SNPs** (A = ancestral allele, D = derived allele (under selection)) (**c**) rs7457618-A/G, AA=AA, AD=AG, DD=GG(**d**) and rs4693058-C/T, AA=CC, AD=CT, DD=TT.

However, both the above-mentioned approaches have limitations. First, associations from the UK biobank have been detected in a different population than Papuans; the transferability of the directionality of the beta values of the associations is therefore limited ^77^. Secondly, we did not find any significant phenotype association for top selection candidate SNPs when correcting for the number of SNPs and phenotypes tested together. That may be because of the low sample size or the choice of documented phenotypes that are not the direct target of selection. Nonetheless, the associations in both analyses with related phenotypes support the hypothesis that cardiovascular phenotypes were a target of selection within PNG highlanders and lowlanders.

### Functional consequences of candidate SNPs

In order to study the potential molecular effects and the most likely target genes of selection candidate SNPs, we investigated their putative regulatory role and impact on the protein structure. Five out of 21 candidate SNPs in PNG highlanders and three out of 23 in PNG lowlanders – including SNPs rs74576183-A/G and rs4693058-C/T whose derived alleles under selection are associated with heart-rate – show significant eQTLs in various GTEx^59^ tissues (Tables S16-S17). Furthermore, 17 out of the 21 putative SNPs driving selection in PNG highlanders and 16 out of 23 in PNG lowlanders are in moderate LD (R2>0.5) with at least one variant with a predicted eQTL in the GETx portal^59^ (Tables S18, S19). Finally, 38 out of the 44 candidate SNPs overlapped with open chromatin regions in at least one epigenome (Figures S5, S6). These results suggest that some of the selection candidate SNPs play a role in gene expression in various primary tissues and cell types.

In addition, we scanned top selected genomic regions for missense variants (Tables S20, S21). We found 191 variants that alter the protein sequence of 18 genes among PNG highlanders selected regions. Regions under selection in PNG lowlanders encompass 85 missense variants that alter 21 genes. In PNG highlanders, one of the regions under selection (chr12:6502552-6612260) overlaps with one missense variant (TAPBPL-G151V), a variant with a exceptionally high derived allele frequency (DAF) in PNG highlanders (DAF = 0.7, <12% in African, Asian or European populations; Table S20). Moreover, this missense variant is in high LD (R2=0.952297) with the candidate SNP, rs74576183-A/G. In contrast, the selection candidate region encompassing GBP overlaps with a missense variant (GBP2-A549P) which is absent in non-Papuan populations and a DAF of 82% in PNG lowlanders (Table S21). This variant is in moderate LD (R2=0.57) with the candidate SNP for the region (rs368120563-T/C). While we expect CLUES top results to be enriched for the causal SNPs of selection, it remains possible that the real targets of selection are SNPs linked to our candidate SNPs. In the case of rs368120563-T/C, we suggest that the linked missense variant GBP2-A549P modifying protein sequence might be the real target of selection for the genomic region.

### Archaic introgressions in loci under selection

We used haplostrips ^55^ to scan regions with selection signatures in PNG highlanders or PNG lowlanders for archaic haplotypes. We observed ten such regions in PNG highlanders (Tables 1, S22). Five of these regions contain archaic SNPs with allele frequencies that are located within the top 10% in Papuans from the SGDP dataset (Table S22). The region with the highest XP-EHH, PBS and Fisher score and carrying *LINC02388* – that might regulate angiogenesis through the HIF/VEGF pathway – carries an archaic haplotype that show high sequence similarity with the Altai Neanderthal. Rs74576183-A/G, the SNP whose derived allele under selection in PNG highlanders is associated with a slower heart rate, is located in a region carrying a Denisovan-like haplotype (Figure S10).

Within regions under selection in PNG lowlanders, we observed six regions with evidence for archaic introgression (Tables 2, S23). Among these is the region encompassing the immunity-related GBP locus (Figure 3) which exhibits the highest selection peak in PNG lowlanders and shows haplotypes with sequence similarities to both Denisovan and Altai Neanderthal. Archaic introgression in this region has previously been reported in Melanesians ^31,35^. But interestingly, the sequence of the introgressed haplotypes does not match with either Vindija ^58^ or Chagyrskaya ^78^ Neanderthals (data not shown). These two Neanderthals are a better reference for the introgressed Neanderthal population in non-African populations than Altai Neanderthal ^58^. This fact and the gene flow between the Altai Neanderthal and Denisova ^57^ would suggest that we most likely observed Denisovan introgression within the GBP locus in the PNG population.

**Figure 3:**
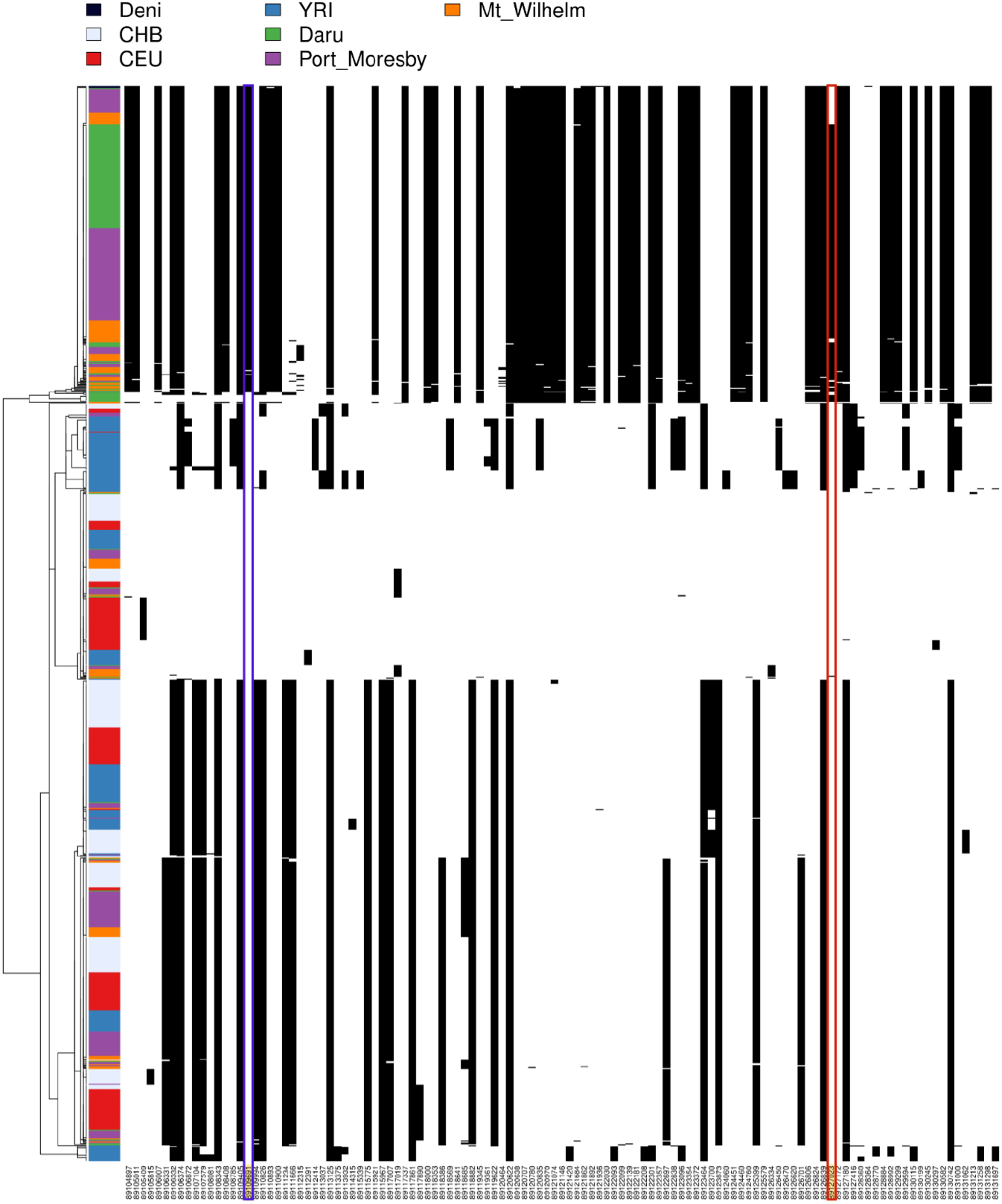
Haplostrips plot for the region chr1:88800562-89326878 overlapping with the GBP locus and under selection in PNG lowlanders. Introgression from Altai Denisovas in PNG for in this region. Derived alleles are plotted in black and ancestral are in white. The introgressed haplotype carry the SNP driving selection for the region (rs368120563-T/C, framed in orange) but the Altai Denisova does not have this particular allele. On the contrary, the missense variant (framed in blue) in LD with rs36812056 is found in the introgressed haplotype and in Denisovan genome

Finally, two candidate SNPs for each studied PNG population (total four SNPs) are exclusively found on introgressed haplotypes (Figure 3, S7-S9) and absent on non-archaic haplotypes. Since these SNPs are not fixed on the archaic haplotypes, this pattern would suggest that the selected mutation appeared after the introgression event and selection of the mutation led to an increase of the introgressed haplotype. Another scenario is that Neandertal and/or Denisovans were variable at this genomic position and introgressed haplotypes with and without the variant and that both types of haplotypes are still segregating in present-day Papuans.

### Cardio Vascular, a target for selection in PNG highlanders

In summary, our analysis of selective pressures in Papuan highlanders suggest that top selected regions encompass genes that might have contributed to counteracting hypoxia detrimental effect in PNG highlanders and that candidate selection SNPs show associations with blood-related phenotypes. For example, the genomic regions on chr12 overlapping with the gene *NCAPD2* demonstrates how hypoxic pressure may have impacted the genome and phenotypes of PNG highlanders. This region shows the third-highest XP-EHH score in PNG highlanders (Table 1, Figure 1). The candidate SNP for this region, rs74576183-A/G (Figure 2), overlaps with the gene *NCAPD2* that is involved in various neurodevelopmental disorders ^79–82^. Similarly, genomic regions under selection in Andeans living at intermediate altitude show enrichment for neuronal-related genes, which might protect their brain from hypoxic damage ^83^. Indeed, hypoxia at altitude impacts brain development and function when exposed during perinatal life ^84,85^ or long after birth ^86,87^. This candidate SNP derived allele under selection shows a significant association with increasing red blood cell count in the UK Biobank (Table S11), and for association with slower heart rate from phenotypes measured in PNG (Figure 2, Table S15) supports adaptation through some cardiovascular related process. The fact that this SNP shows significant eQTL associations and overlaps with open chromatin state in multiple tissues would supports its role in gene expression regulation. However, because this SNPs is in high LD with a missense variant with high DAF in PNG Highlanders but rare in other populations (Table S20), it is also possible that the real target for selection might be the missense variant (TAPBPL-G151V) that leads to changes in the TAPBPL protein that is associated with antigen processing. This region under selection overlap with Denisovan-like archaic haplotypes (Tables 1, S22, Figure S10) but neither the candidate SNP nor the missense variant derived allele are found in PNG individuals that carry this archaic haplotype (Figure S10).

### Immunity, a target for selection in PNG lowlanders

Similarly, the region containing the gene *SEC31A* and rs4693058-C/T, the candidate SNP for this region (Figure 2), are of particular interest to selection for pathogenic pressure in PNG lowlanders. Indeed *SEC31A* ^73^ might play a role in immune processes, and the derived allele under selection of rs4693058-C/T, the candidate SNP for this locus, shows a significant association with various white cells percentages and counts (Table S13). Interestingly derived allele T under selection of rs4693058-C/T shows a suggestive association with faster heart rate (Figure 2). But once again, we suggest that heart rate might be a proxy for other phenotypes (here the white cells count ^88^). Because rs4693058-C/T show significant eQTLs and overlaps with open chromatin states in multiple tissues (Table S17, Figure S6), we hypothesise that it impacts gene expression regulation. This region under selection overlaps with an introgressed haplotype from Denisovan, but the introgressed haplotype does not carry the derived allele of the candidate SNP (Figure S11).

Finally, the regions with the highest XP-EHH, PBS and Fisher Score in PNG lowlanders (Figure 1, Tables 2, S6), includes several genes from the guanine-binding protein (GBP) associated with immunity to diverse pathogens ^76^. Especially, Apinjoh et al. reported an association between *GBP7* variant and higher malaria symptoms in the Cameroon population ^89^, suggesting this region might be selected due to malaria. The candidate SNP, rs368120563-T/C, is in LD with a missense variant (GBP2-A549P) with a high DAF in PNG lowlanders (DAF=0.82) but absent in non-Papuan populations (Table S21). This missense variant is part of the top 5 SNPs given by CLUES for the region (Table S10). That might suggest that we failed to identify the real selection driving SNP when limiting the candidate SNPs to the first top one. This particular missense variant might be the causal SNP and selection might have targeted a change in the GBP2 protein sequence. This GBP locus carries a Denisovan-like haplotype that includes both the candidate variant of the region (rs368120563-T/C) and the missense variant (GBP2-A549P) in PNG populations. Moreover, the missense variant can be found in the Denisovan genome, but the candidate SNP is not present in the Denisovan or any of the high coverage Neandertal genomes (Figure 3). That pattern is compatible with the scenario where the candidate variant appeared after the introgression and that the introgressed haplotype frequency increased in the PNG populations driven by the selection acting on this variant. The alternative hypothesis would be that the candidate variant is not the target of selection (most likely the missense variant is), and the candidate variant is hitchhiked with the selected and introgressed haplotype.

## Conclusion

In this paper we investigated selection in PNG highlanders and PNG lowlanders and detected 21 and 23 genomic regions under positive selection, respectively. Within each candidate selection region, we identified the SNP that most likely drives selection and explore their association with several phenotypes measured within our dataset or UK Biobank summary statistics. The genes in regions that show selection signals in PNG highlanders are associated with HIF pathway regulation, brain development, blood composition and immunity. PNG lowlanders show selection for immune system. In both populations, one of the candidate SNPs suggests an association with heart rate. This SNP and several top SNPs were also significantly associated with several blood composition phenotypes in the UK Biobank. Further studies will be needed to clarify the complexity of the PNG’s haematological responses to hypoxia and pathogenic pressures. We found that 16 regions under selection −10 in PNG highlanders and 6 in PNG lowlanders – carry archaic introgression. Out of which, two candidate SNPs from both populations (a total of four) reside directly inside the introgressed haplotypes suggesting adaptive introgression. Our results suggest that selection in PNG highlanders and lowlanders was partially targetting introgressed haplotypes from Neandertals and Densiovans. This study demonstrates that both PNG highlanders and PNG lowlanders carry signatures of positive selection and that the associated phenotypes largely match with the challenges they faced due to the environmental differences.

## Authors contribution

F.-X.R., N.B., M.L., T.O. and M.Me. designed the study. F.-X.R, N.B., M.L., J.K., N.E. and J.M. collected the data. V.M., A.B., and J.F.D. generated whole-genome sequences. M.A., N.B., G.H., V.P., D.Y., R.K. and M.Mo. performed the data analysis. F.-X.R., M.Me. and M.P.C. provided resources and logistics. M.A., N.B., M.Mo. and F-X.R. wrote the manuscript with the contribution from all the co-authors.

## Data availability

PNG highlanders (n=38) and lowlanders (n=46) sequenced genomes are on the European Genome-Phenome data repository: EGAXXX.

## Funding

M.A. was supported by the European Union through the European Regional Development Fund (Project No. 2014-2020.4.01.16-0030)., G.H., V.P., R.K., M.D., T.O. and M.Mo. were supported by the European Union through Horizon 2020 research and innovation programme under grant no 810645 and the European Regional Development Fund project no. MOBEC008. This work was supported by the French Ministry of Foreign and European Affairs (https://www.diplomatie.gouv.fr) (French Prehistoric Mission in Papua New Guinea to F.-X.R.), the French Embassy in Papua New Guinea (https://pg.ambafrance.org), and the University of Papua New Guinea, Archaeology Laboratory Group. We acknowledge support from the LabEx TULIP, France (https://www.labex-tulip.fr) (to F.-X. R. and N. B.) and the French Ministry of Research grant Agence Nationale de la Recherche (https://anr.fr) number ANR-20-CE12-0003-01 (to F.-X.R.); from the Leakey Foundation (https://leakeyfoundation.org) (to N. B.). The CNRGH sequencing platform was supported by the “France Génomique” national infrastructure, funded as part of the “Investissements d’Avenir” program managed by the “Agence Nationale pour la Recherche” (contract ANR-10-INBS-09).

## Acknowledgments

We kindly thank F.-X. Ricaut for giving us access to additional PNG whole genome sequences from Daru, Mt. Wilhelm and Port Moresby. These data were generated at the National Center of Human Genomics Research (France) or the KCCG Sequencing Laboratory (Garvan Institute of Medical Research, Australia). We thank Ray Tobler (Australian National University), Roxanne Tsang (Centre for Social and Cultural Research, Griffith University, Australia), Kylie Sesuki and Teppsy Beni (School of Humanities and Social Sciences, University of Papua New Guinea), and Alois Kuaso and Kenneth Miamba (National Museum and Art Gallery, Papua New Guinea) for their help during the sampling campaigns. We especially thank all of our study participants. Data analyses were carried out in part in the High-Performance Computing Center of the University of Tartu.

## Competing interest

The authors declare no competing interest.

## Notes

### Competing Interest Statement

The authors have declared no competing interest.

