## Supplementary Figures and Notes for "Positive selection in the genomes of two Papua New Guinean populations at distinct altitude levels"

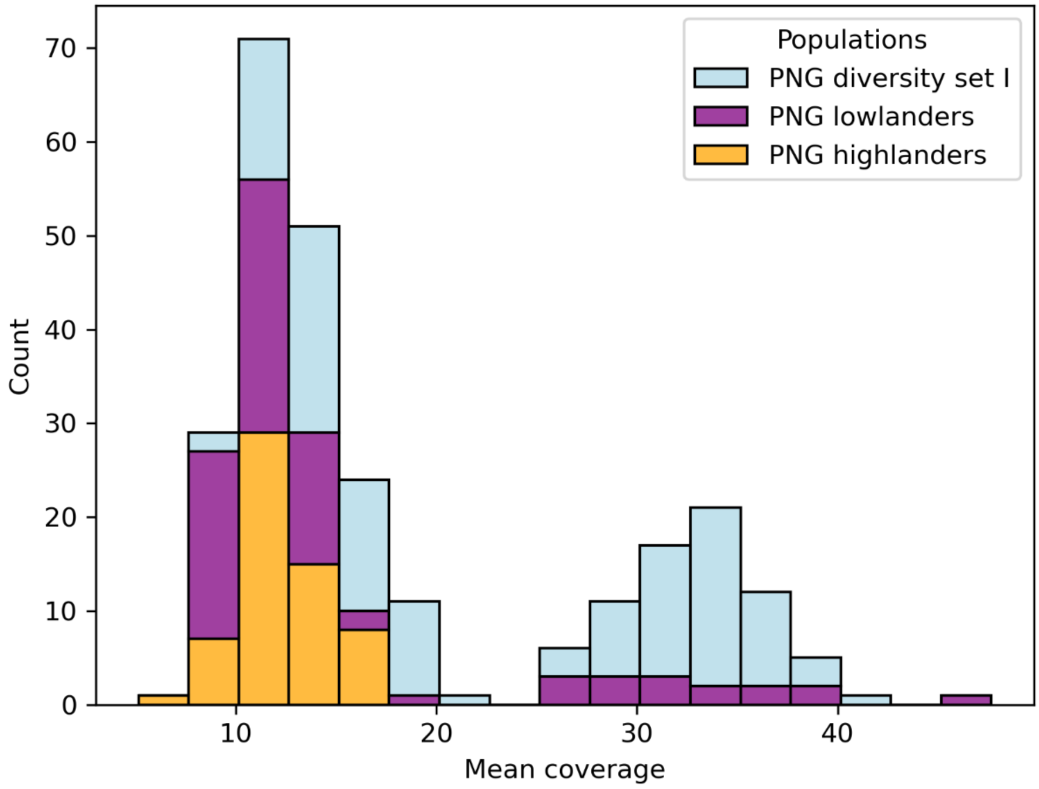

**Figure S1:** Distribution of mean coverage within the autosomes interval in Port Moresby (PNG diversity set I), PNG highlanders (Mt Wilhelm) and PNG lowlanders (Daru) sequences

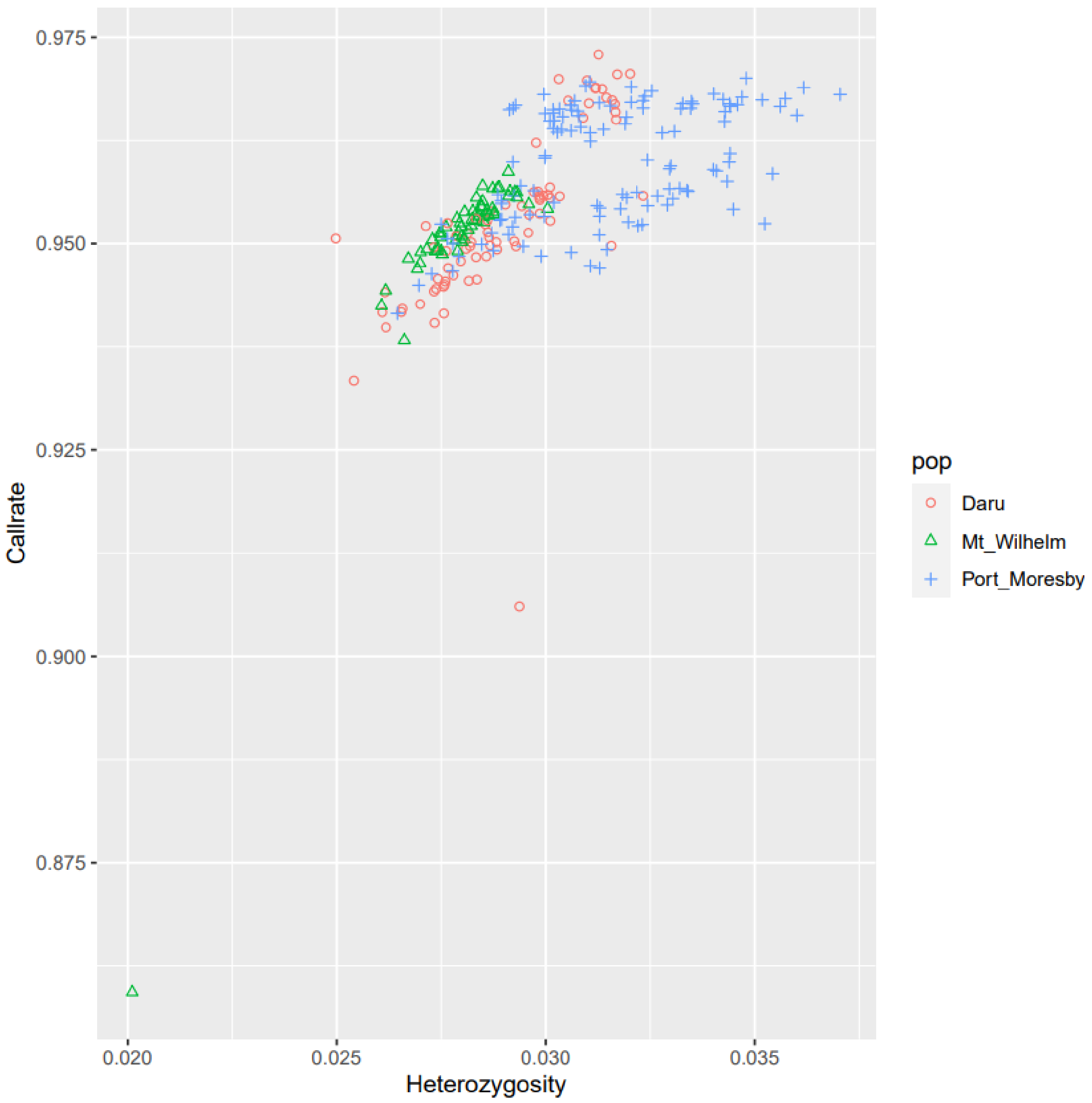

**Figure S2:** Heterozygosity to callrate in within autosome interval in PNG diversity set I (Port Moresby), PNG highlanders(Mt Wilhelm), PNG lowlanders (Daru) sequences

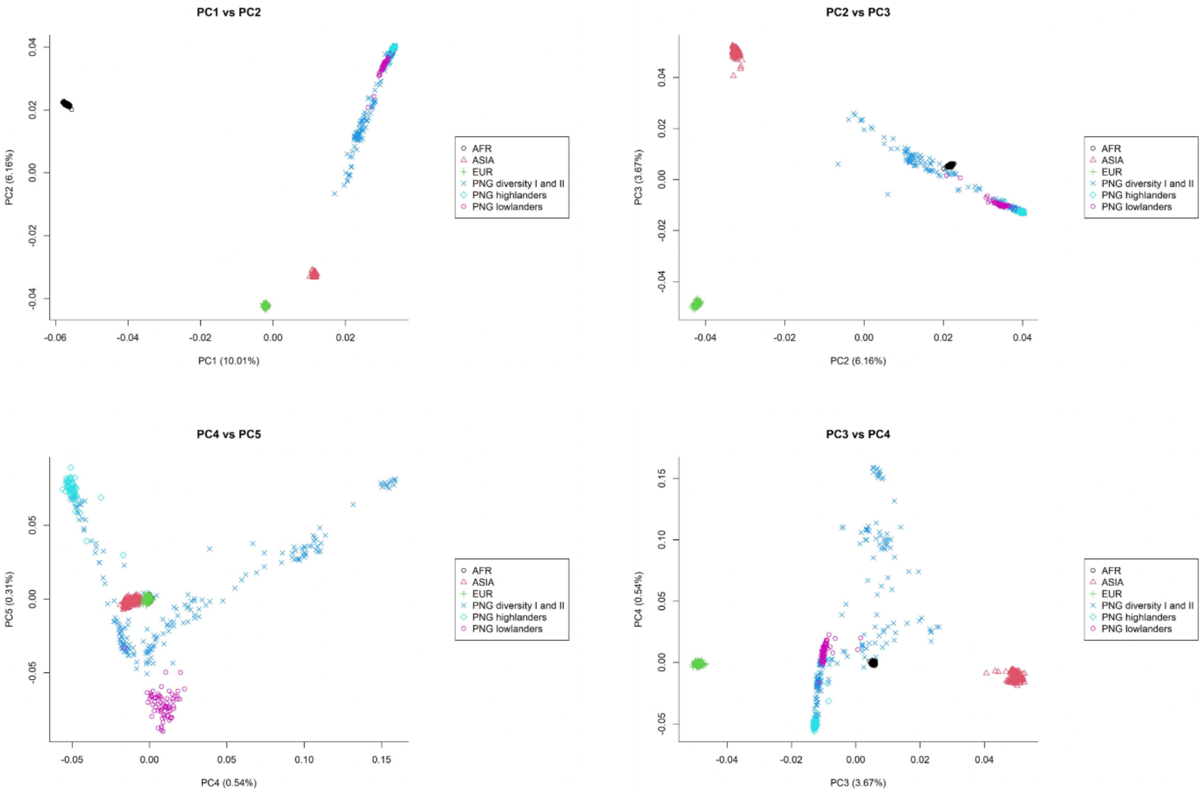

**Figure S3:** PCA plots fromPC1 to PC5. AFR=YRI and ESN from1000G, ASIA=CHB and KHV from1000G, EUR= GBR and CEU from 1000G, PNG diversity I and II II [1–6].

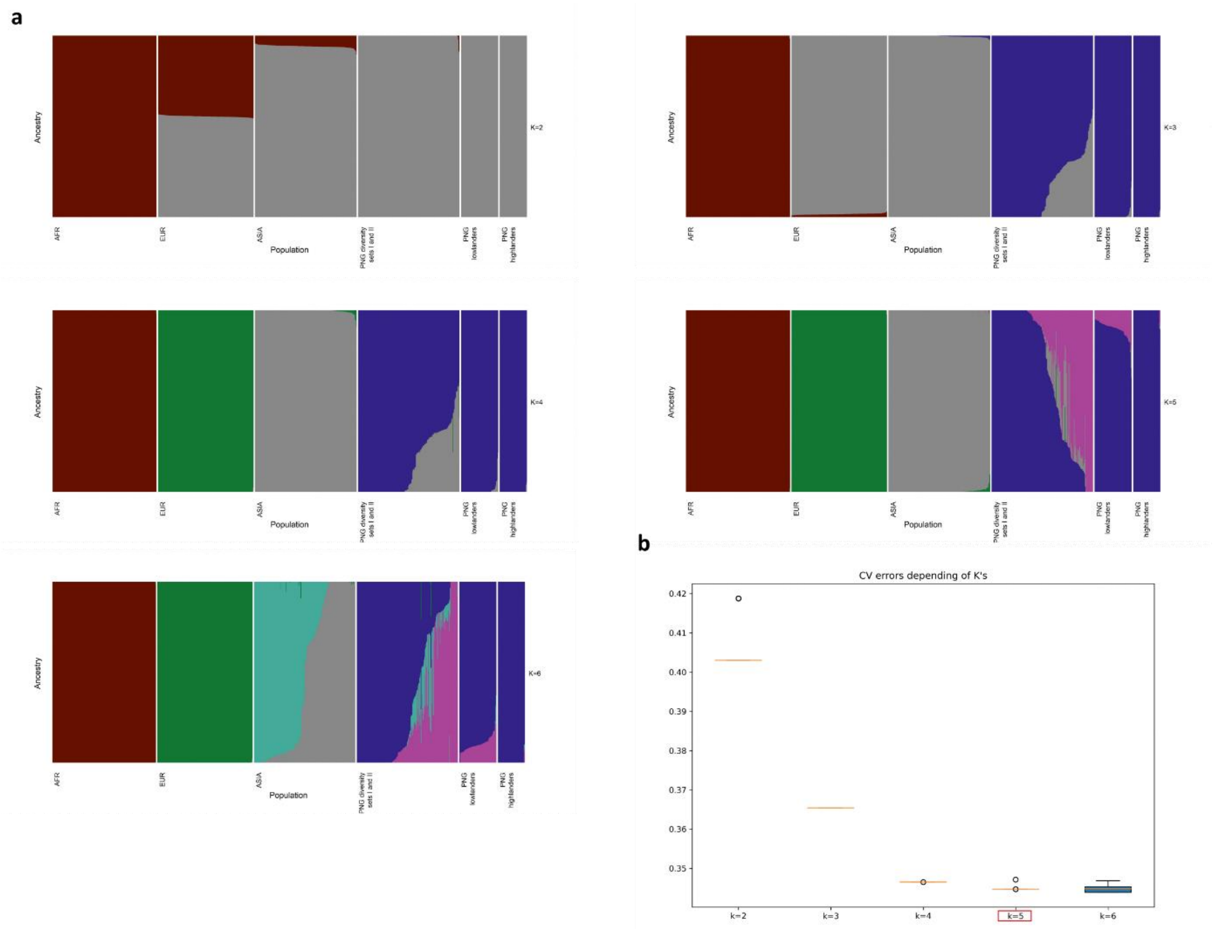

**Figure S4: a ADMIXTURE plots from K=2 to K=8**. AFR=YRI and ESN from 1000G, ASIA=CHB and KHV from 1000G, EUR= GBR and CEU from1000G, PNG diversity I and II [1–6] **b. Cross Validation errors for ADMIXTURE.** CV errors were computed for 50 independent runs of the ADMIXTURE analysis from K=2 to K=6. K=5 has the lowest CV error (framed in red).

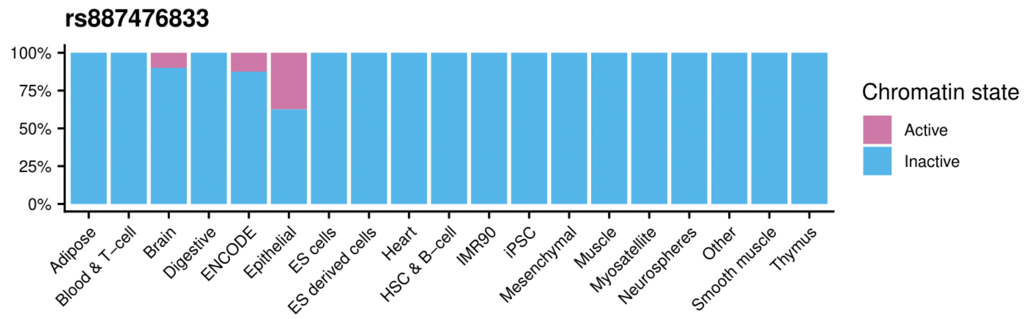
rs887476833
100%
75%
50%
25%
0%
Adipose
Blood & T−cell
Brain
Digestive
ENCODE
Epithelial
ES cells
ES derived cells
Heart
HSC & B−cell
IMR90
iPSC
Mesenchymal
Muscle
Myosatellite
Neurospheres
Other
Smooth muscle
Thymus
Chromatin state
Active
Inactive

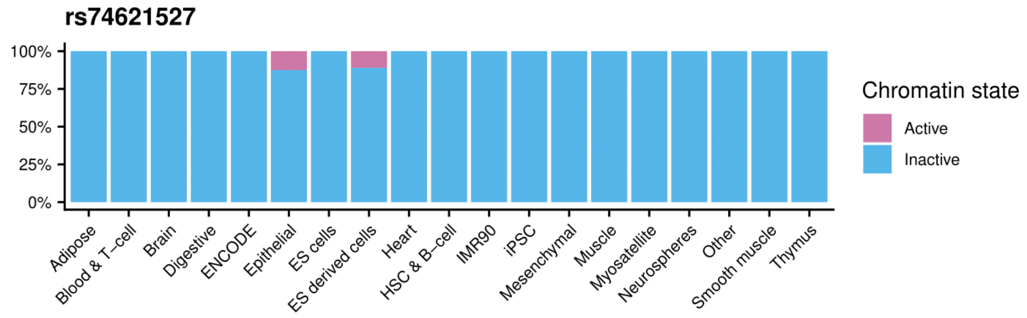
rs74621527
100%
75%
50%
25%
0%
Adipose
Blood & T−cell
Brain
Digestive
ENCODE
Epithelial
ES cells
ES derived cells
Heart
HSC & B−cell
IMR90
iPSC
Mesenchymal
Muscle
Myosatellite
Neurospheres
Other
Smooth muscle
Thymus
Chromatin state
Active
Inactive

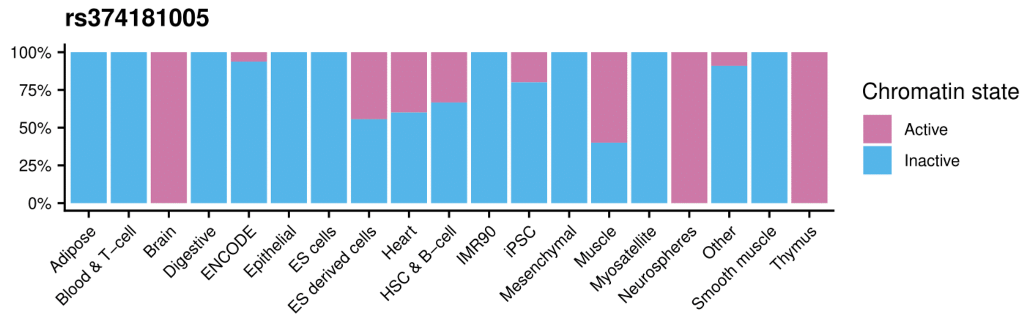
rs374181005
100%
75%
50%
25%
0%
Adipose
Blood & T−cell
Brain
Digestive
ENCODE
Epithelial
ES cells
ES derived cells
Heart
HSC & B−cell
IMR90
iPSC
Mesenchymal
Muscle
Myosatellite
Neurospheres
Other
Smooth muscle
Thymus
Chromatin state
Active
Inactive

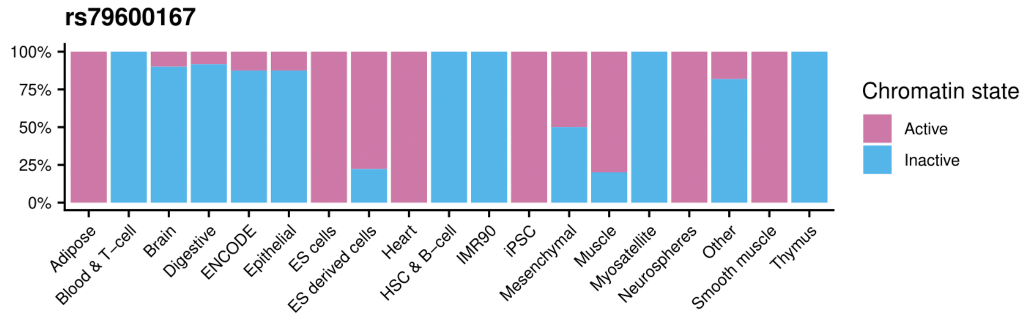
rs79600167
100%
75%
50%
25%
0%
Adipose
Blood & T−cell
Brain
Digestive
ENCODE
Epithelial
ES cells
ES derived cells
Heart
HSC & B−cell
IMR90
iPSC
Mesenchymal
Muscle
Myosatellite
Neurospheres
Other
Smooth muscle
Thymus
Chromatin state
Active
Inactive

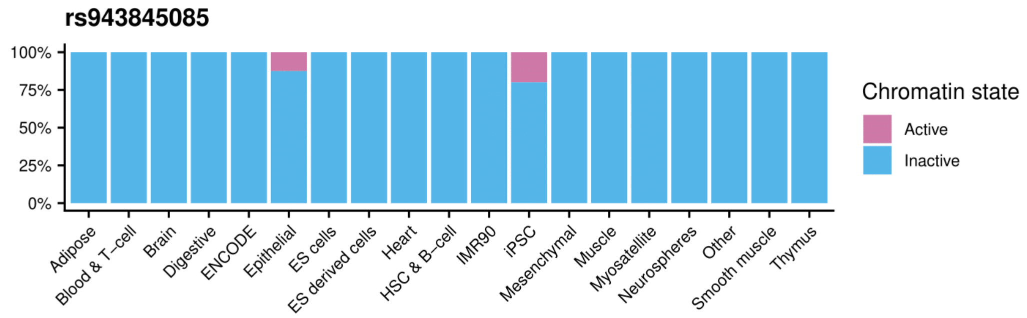
rs943845085
100%
75%
50%
25%
0%
Adipose
Blood & T−cell
Brain
Digestive
ENCODE
Epithelial
ES cells
ES derived cells
Heart
HSC & B−cell
IMR90
iPSC
Mesenchymal
Muscle
Myosatellite
Neurospheres
Other
Smooth muscle
Thymus
Chromatin state
Active
Inactive

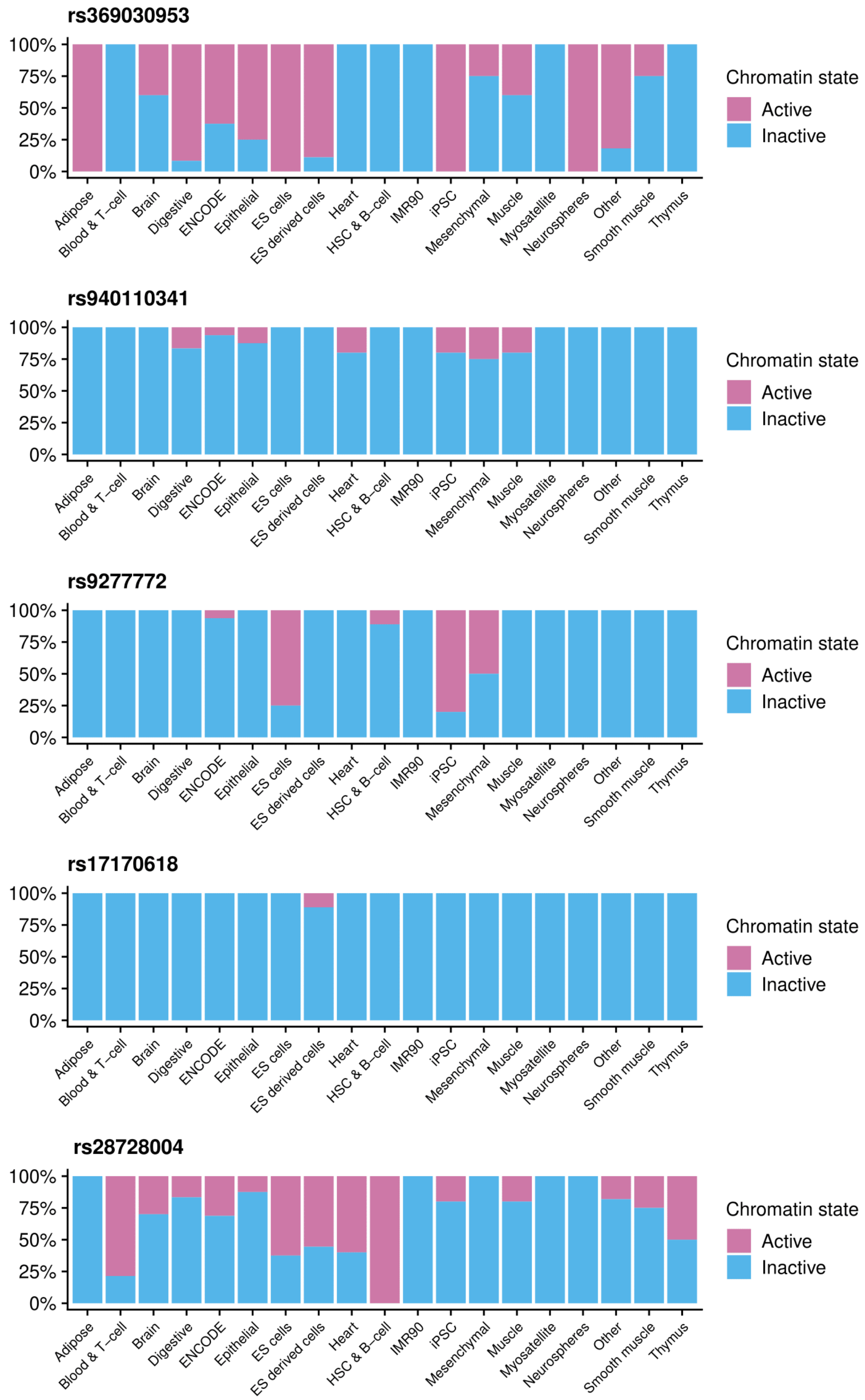
rs369030953
rs940110341
rs9277772
rs17170618
rs28728004
100%
75%
50%
25%
0%
Adipose
Blood & T−cell
Brain
Digestive
ENCODE
Epithelial
ES cells
ES derived cells
Heart
HSC & B−cell
IMR90
iPSC
Mesenchymal
Muscle
Myosatellite
Neurospheres
Other
Smooth muscle
Thymus
Chromatin state
Active
Inactive

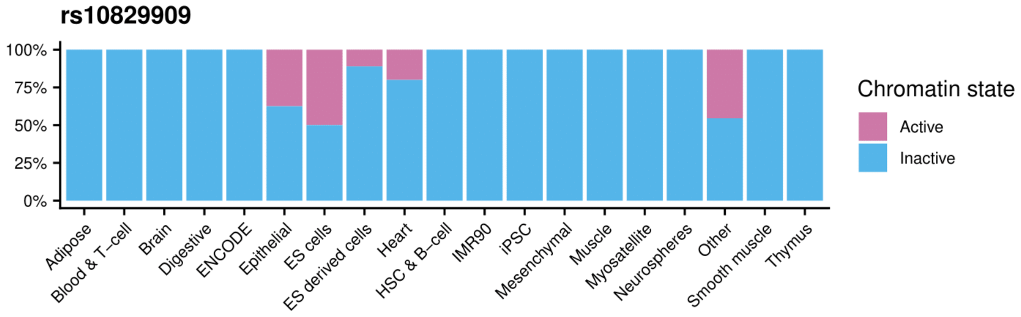
rs10829909
100%
75%
50%
25%
0%
Adipose
Blood & T−cell
Brain
Digestive
ENCODE
Epithelial
ES cells
ES derived cells
Heart
HSC & B−cell
IMR90
iPSC
Mesenchymal
Muscle
Myosatellite
Neurospheres
Other
Smooth muscle
Thymus
Chromatin state
Active
Inactive

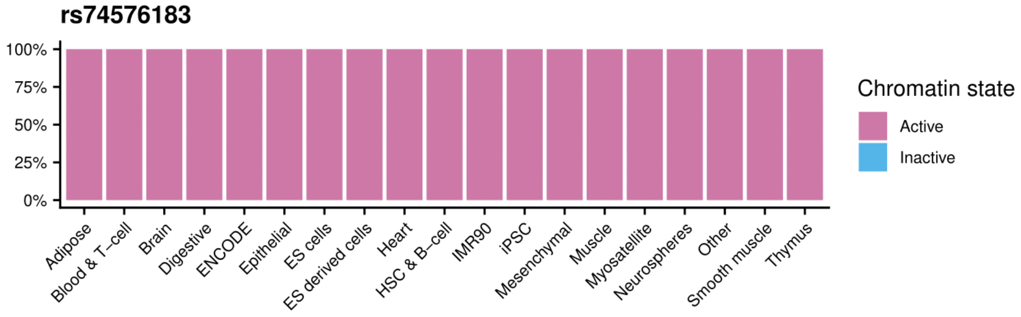
rs74576183
100%
75%
50%
25%
0%
Adipose
Blood & T−cell
Brain
Digestive
ENCODE
Epithelial
ES cells
ES derived cells
Heart
HSC & B−cell
IMR90
iPSC
Mesenchymal
Muscle
Myosatellite
Neurospheres
Other
Smooth muscle
Thymus
Chromatin state
Active
Inactive

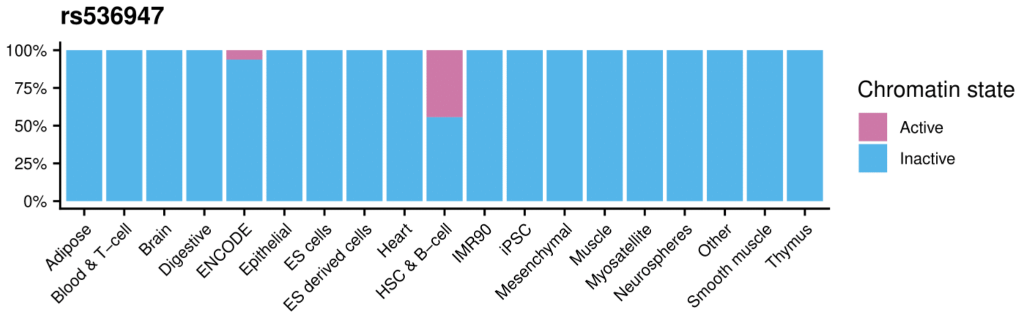
rs536947
100%
75%
50%
25%
0%
Adipose
Blood & T−cell
Brain
Digestive
ENCODE
Epithelial
ES cells
ES derived cells
Heart
HSC & B−cell
IMR90
iPSC
Mesenchymal
Muscle
Myosatellite
Neurospheres
Other
Smooth muscle
Thymus
Chromatin state
Active
Inactive

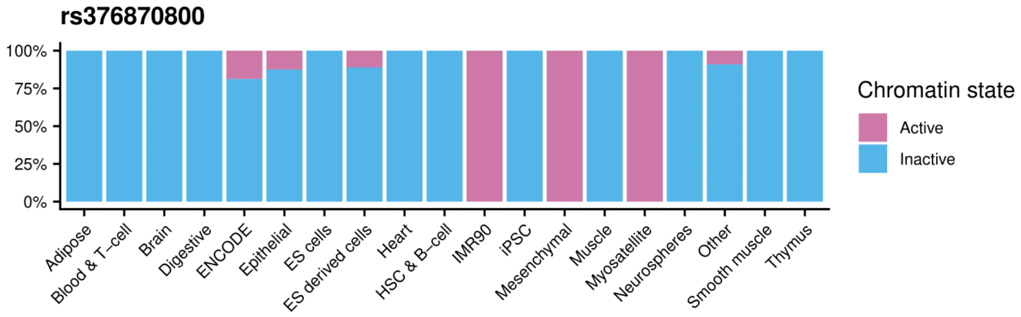
rs376870800
100%
75%
50%
25%
0%
Adipose
Blood & T−cell
Brain
Digestive
ENCODE
Epithelial
ES cells
ES derived cells
Heart
HSC & B−cell
IMR90
iPSC
Mesenchymal
Muscle
Myosatellite
Neurospheres
Other
Smooth muscle
Thymus
Chromatin state
Active
Inactive

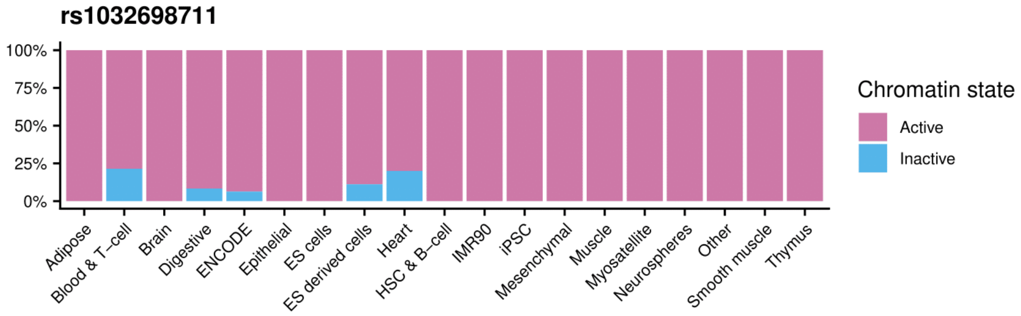
rs1032698711
100%
75%
50%
25%
0%
Adipose
Blood & T−cell
Brain
Digestive
ENCODE
Epithelial
ES cells
ES derived cells
Heart
HSC & B−cell
IMR90
iPSC
Mesenchymal
Muscle
Myosatellite
Neurospheres
Other
Smooth muscle
Thymus
Chromatin state
Active
Inactive

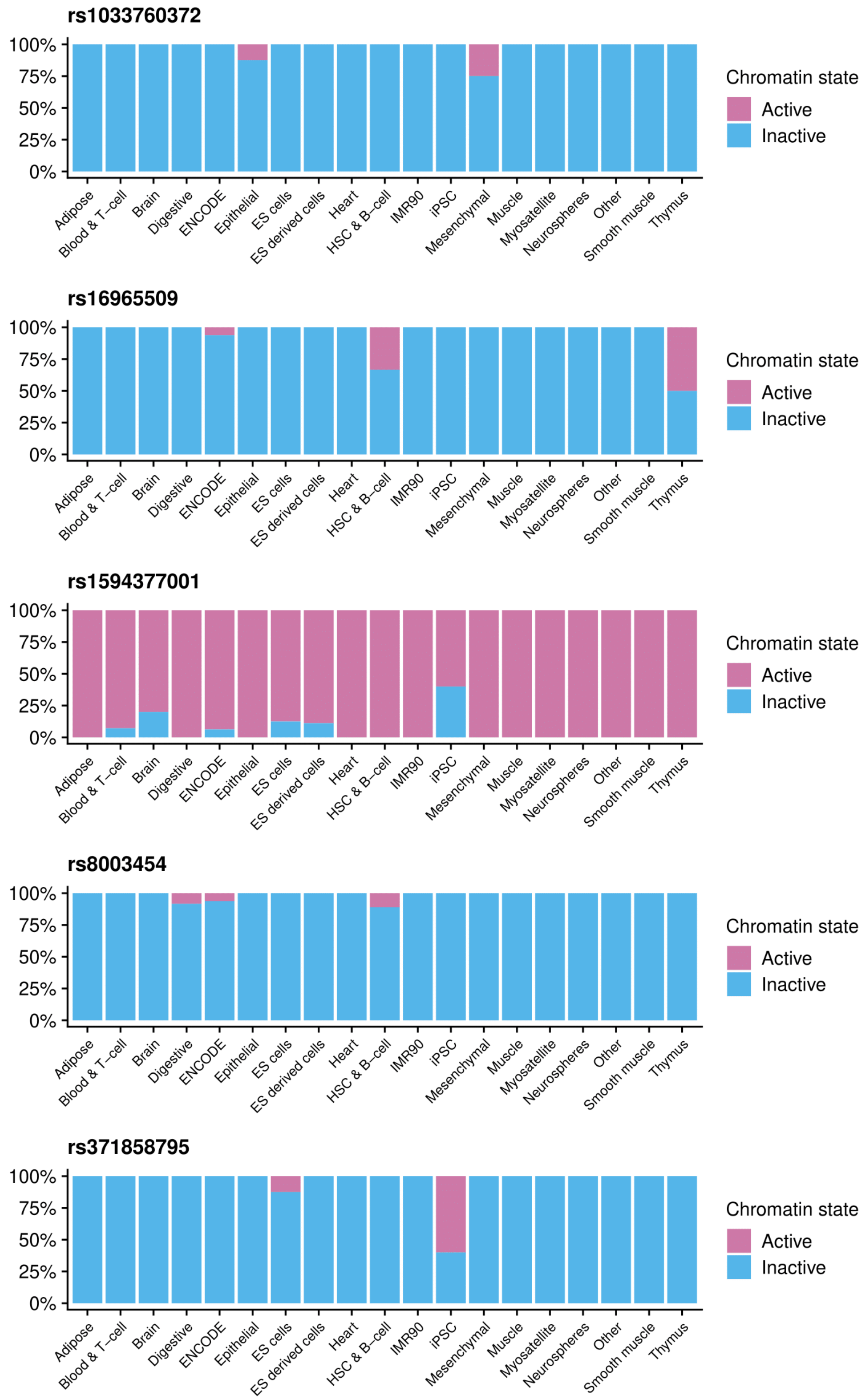
rs1033760372
100%
75%
50%
25%
0%
Adipose
Blood & T–cell
Brain
Digestive
ENCODE
Epithelial
ES cells
ES derived cells
Heart
HSC & B–cell
IMR90
iPSC
Mesenchymal
Muscle
Myosatellite
Neurospheres
Other
Smooth muscle
Thymus
Chromatin state
Active
Inactive
rs16965509
100%
75%
50%
25%
0%
Adipose
Blood & T–cell
Brain
Digestive
ENCODE
Epithelial
ES cells
ES derived cells
Heart
HSC & B–cell
IMR90
iPSC
Mesenchymal
Muscle
Myosatellite
Neurospheres
Other
Smooth muscle
Thymus
Chromatin state
Active
Inactive
rs1594377001
100%
75%
50%
25%
0%
Adipose
Blood & T–cell
Brain
Digestive
ENCODE
Epithelial
ES cells
ES derived cells
Heart
HSC & B–cell
IMR90
iPSC
Mesenchymal
Muscle
Myosatellite
Neurospheres
Other
Smooth muscle
Thymus
Chromatin state
Active
Inactive
rs8003454
100%
75%
50%
25%
0%
Adipose
Blood & T–cell
Brain
Digestive
ENCODE
Epithelial
ES cells
ES derived cells
Heart
HSC & B–cell
IMR90
iPSC
Mesenchymal
Muscle
Myosatellite
Neurospheres
Other
Smooth muscle
Thymus
Chromatin state
Active
Inactive
rs371858795
100%
75%
50%
25%
0%
Adipose
Blood & T–cell
Brain
Digestive
ENCODE
Epithelial
ES cells
ES derived cells
Heart
HSC & B–cell
IMR90
iPSC
Mesenchymal
Muscle
Myosatellite
Neurospheres
Other
Smooth muscle
Thymus
Chromatin state
Active
Inactive

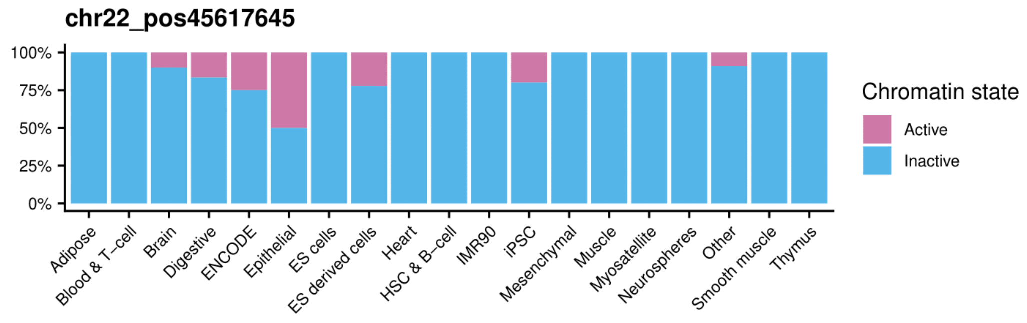

**Figure S5:** Chromatin state of the candidate SNPs of PNG highlanders for the epigenomes of the Roadmap project clusters in tissues

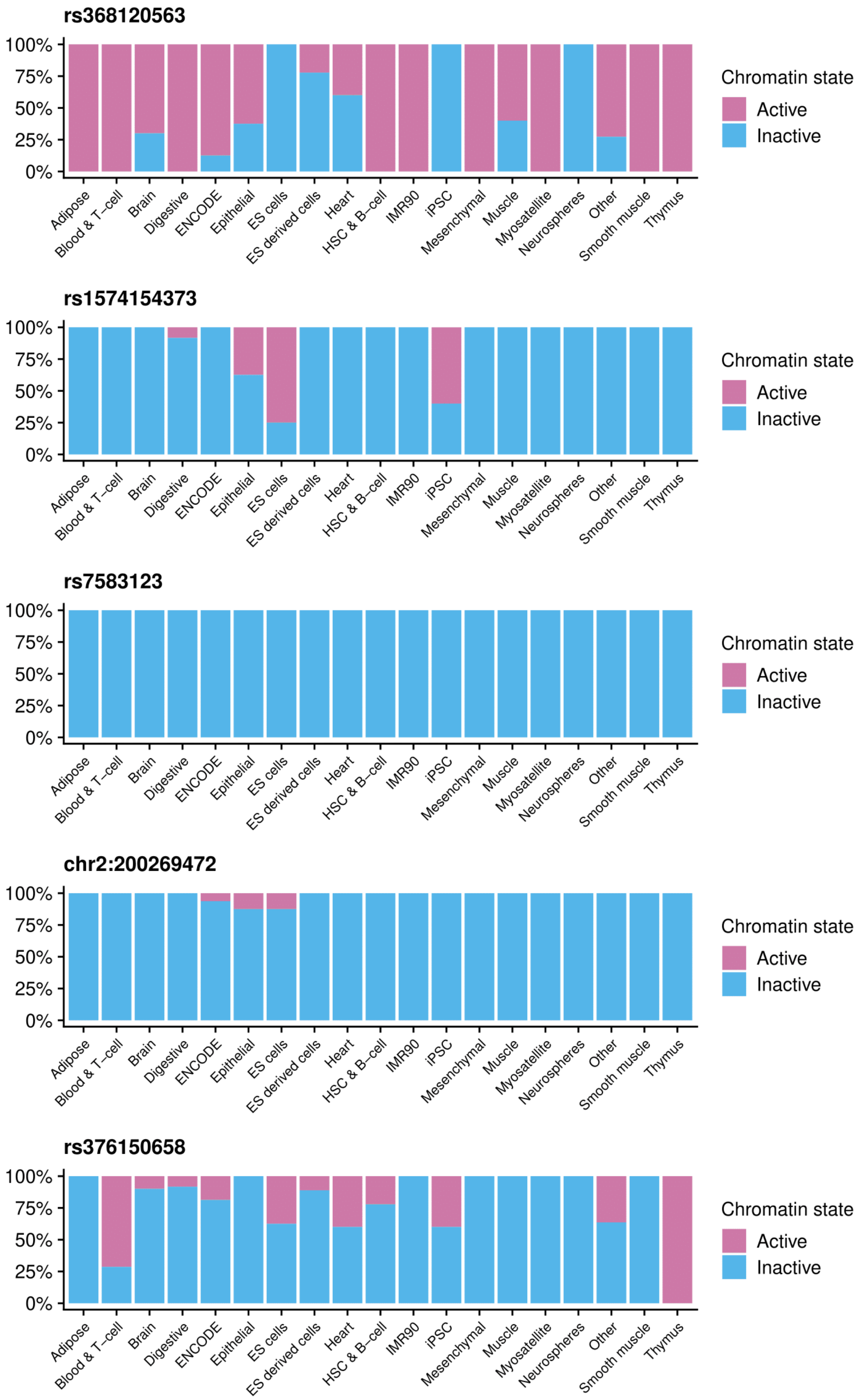
rs368120563
rs1574154373
rs7583123
chr2:200269472
rs376150658
100%
75%
50%
25%
0%
Adipose
Blood & T−cell
Brain
Digestive
ENCODE
Epithelial
ES cells
ES derived cells
Heart
HSC & B−cell
IMR90
iPSC
Mesenchymal
Muscle
Myosatellite
Neurospheres
Other
Smooth muscle
Thymus
Chromatin state
Active
Inactive

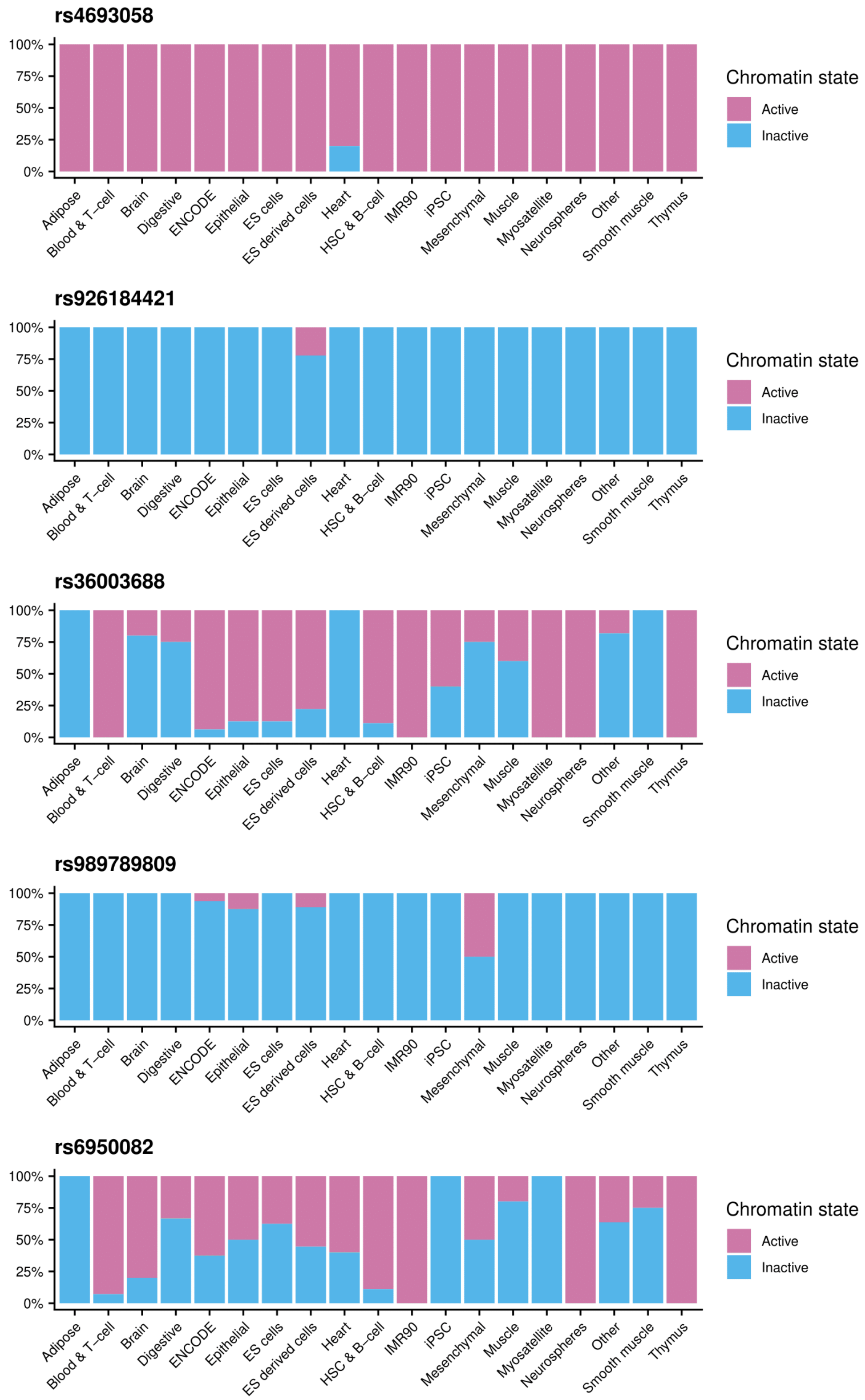
rs4693058
100%
75%
50%
25%
0%
Adipose
Blood & T−cell
Brain
Digestive
ENCODE
Epithelial
ES cells
ES derived cells
Heart
HSC & B−cell
IMR90
iPSC
Mesenchymal
Muscle
Myosatellite
Neurospheres
Other
Smooth muscle
Thymus
Chromatin state
Active
Inactive
rs926184421
100%
75%
50%
25%
0%
Adipose
Blood & T−cell
Brain
Digestive
ENCODE
Epithelial
ES cells
ES derived cells
Heart
HSC & B−cell
IMR90
iPSC
Mesenchymal
Muscle
Myosatellite
Neurospheres
Other
Smooth muscle
Thymus
Chromatin state
Active
Inactive
rs36003688
100%
75%
50%
25%
0%
Adipose
Blood & T−cell
Brain
Digestive
ENCODE
Epithelial
ES cells
ES derived cells
Heart
HSC & B−cell
IMR90
iPSC
Mesenchymal
Muscle
Myosatellite
Neurospheres
Other
Smooth muscle
Thymus
Chromatin state
Active
Inactive
rs989789809
100%
75%
50%
25%
0%
Adipose
Blood & T−cell
Brain
Digestive
ENCODE
Epithelial
ES cells
ES derived cells
Heart
HSC & B−cell
IMR90
iPSC
Mesenchymal
Muscle
Myosatellite
Neurospheres
Other
Smooth muscle
Thymus
Chromatin state
Active
Inactive
rs6950082
100%
75%
50%
25%
0%
Adipose
Blood & T−cell
Brain
Digestive
ENCODE
Epithelial
ES cells
ES derived cells
Heart
HSC & B−cell
IMR90
iPSC
Mesenchymal
Muscle
Myosatellite
Neurospheres
Other
Smooth muscle
Thymus
Chromatin state
Active
Inactive

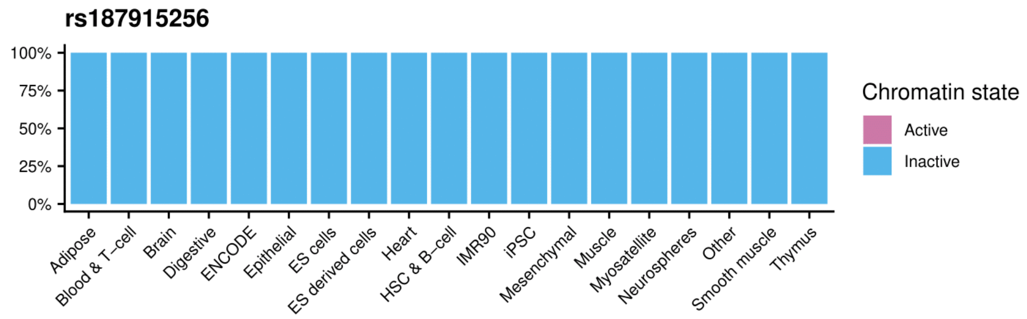
rs187915256
100%
75%
50%
25%
0%
Adipose
Blood & T-cell
Brain
Digestive
ENCODE
Epithelial
ES cells
ES derived cells
Heart
HSC & B-cell
IMR90
iPSC
Mesenchymal
Muscle
Myosatellite
Neurospheres
Other
Smooth muscle
Thymus
Chromatin state
Active
Inactive

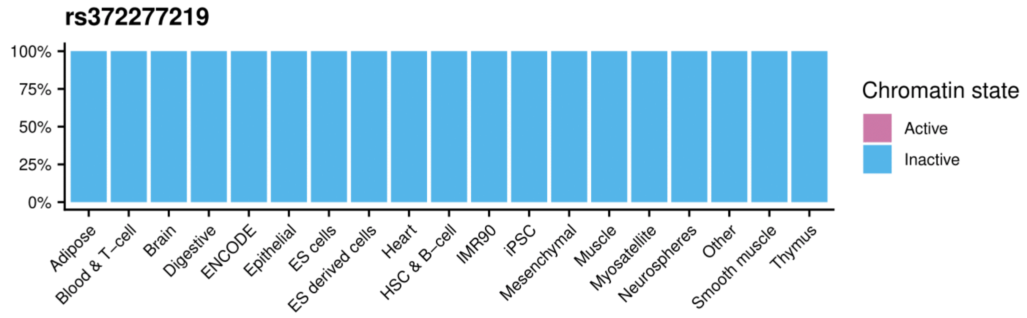
rs372277219
100%
75%
50%
25%
0%
Adipose
Blood & T-cell
Brain
Digestive
ENCODE
Epithelial
ES cells
ES derived cells
Heart
HSC & B-cell
IMR90
iPSC
Mesenchymal
Muscle
Myosatellite
Neurospheres
Other
Smooth muscle
Thymus
Chromatin state
Active
Inactive

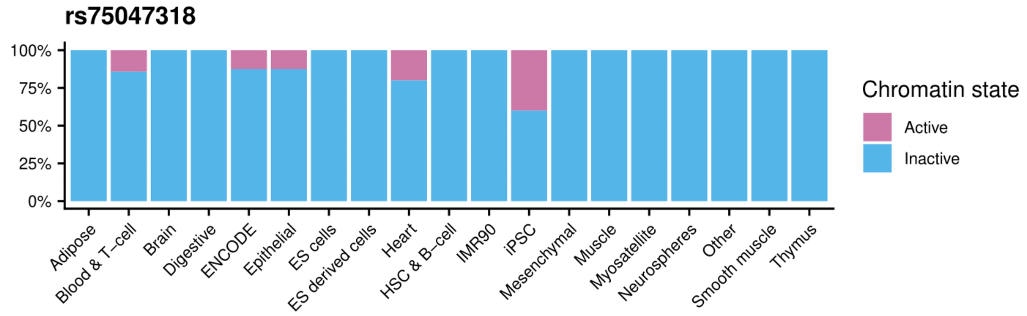
rs75047318
100%
75%
50%
25%
0%
Adipose
Blood & T-cell
Brain
Digestive
ENCODE
Epithelial
ES cells
ES derived cells
Heart
HSC & B-cell
IMR90
iPSC
Mesenchymal
Muscle
Myosatellite
Neurospheres
Other
Smooth muscle
Thymus
Chromatin state
Active
Inactive

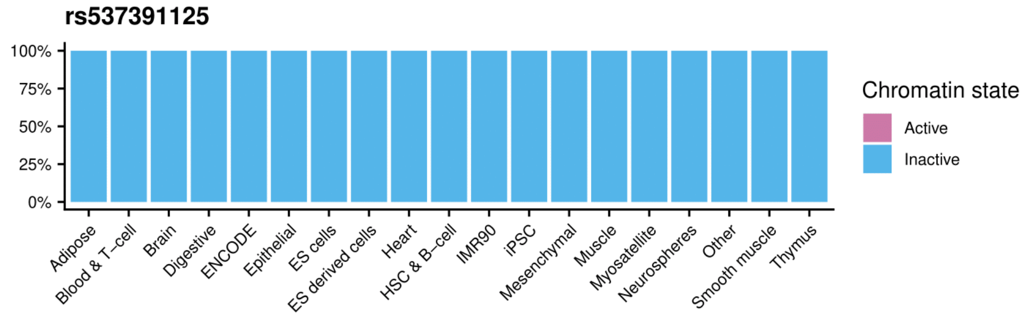
rs537391125
100%
75%
50%
25%
0%
Adipose
Blood & T-cell
Brain
Digestive
ENCODE
Epithelial
ES cells
ES derived cells
Heart
HSC & B-cell
IMR90
iPSC
Mesenchymal
Muscle
Myosatellite
Neurospheres
Other
Smooth muscle
Thymus
Chromatin state
Active
Inactive

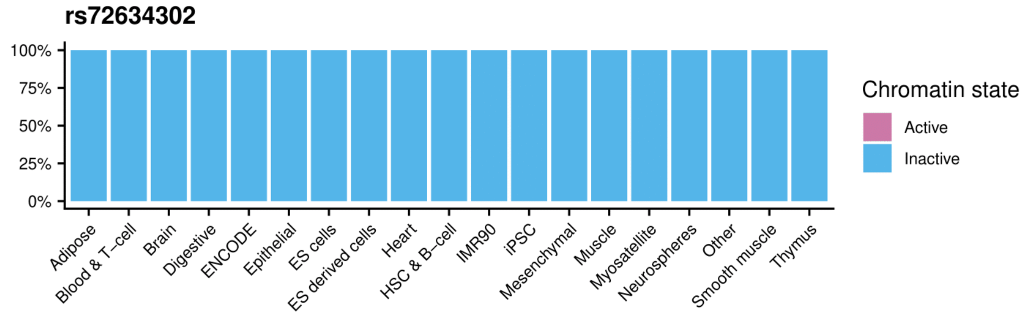
rs72634302
100%
75%
50%
25%
0%
Adipose
Blood & T-cell
Brain
Digestive
ENCODE
Epithelial
ES cells
ES derived cells
Heart
HSC & B-cell
IMR90
iPSC
Mesenchymal
Muscle
Myosatellite
Neurospheres
Other
Smooth muscle
Thymus
Chromatin state
Active
Inactive

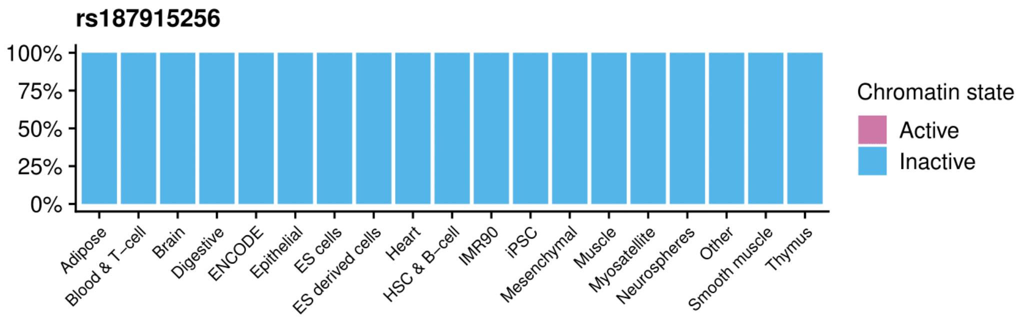
rs187915256
100%
75%
50%
25%
0%
Adipose
Blood & T–cell
Brain
Digestive
ENCODE
Epithelial
ES cells
ES derived cells
Heart
HSC & B–cell
IMR90
iPSC
Mesenchymal
Muscle
Myosatellite
Neurospheres
Other
Smooth muscle
Thymus
Chromatin state
Active
Inactive

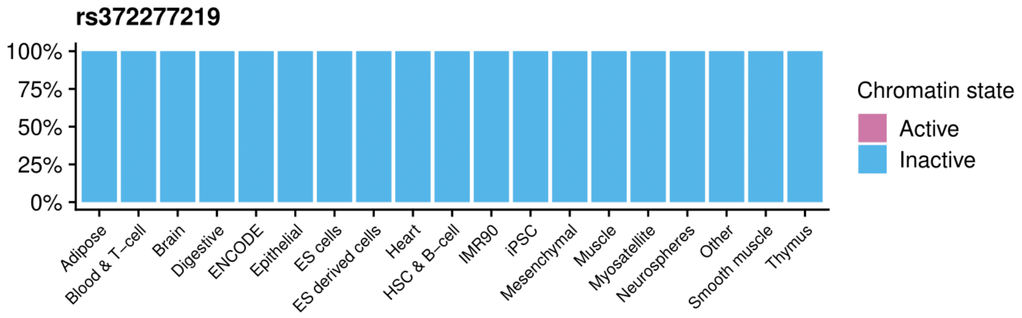
rs372277219
100%
75%
50%
25%
0%
Adipose
Blood & T–cell
Brain
Digestive
ENCODE
Epithelial
ES cells
ES derived cells
Heart
HSC & B–cell
IMR90
iPSC
Mesenchymal
Muscle
Myosatellite
Neurospheres
Other
Smooth muscle
Thymus
Chromatin state
Active
Inactive

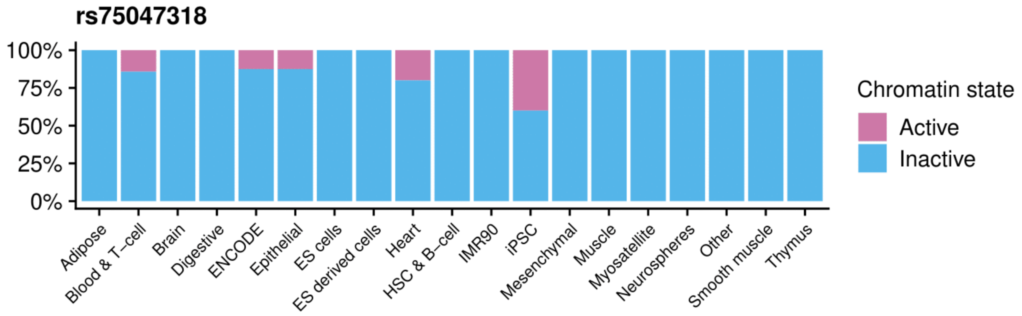
rs75047318
100%
75%
50%
25%
0%
Adipose
Blood & T–cell
Brain
Digestive
ENCODE
Epithelial
ES cells
ES derived cells
Heart
HSC & B–cell
IMR90
iPSC
Mesenchymal
Muscle
Myosatellite
Neurospheres
Other
Smooth muscle
Thymus
Chromatin state
Active
Inactive

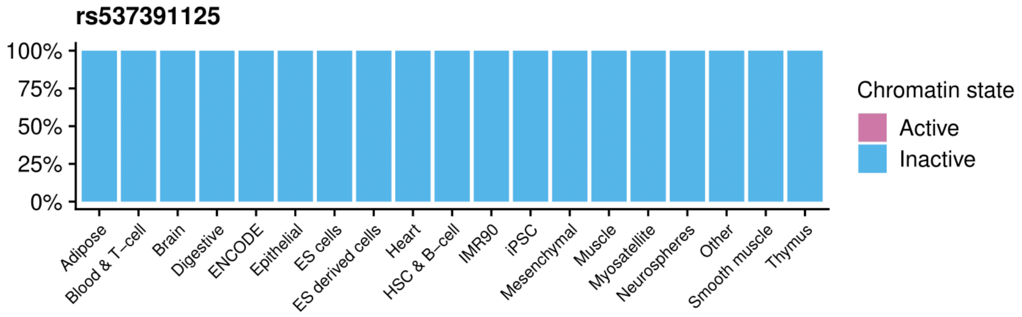
rs537391125
100%
75%
50%
25%
0%
Adipose
Blood & T–cell
Brain
Digestive
ENCODE
Epithelial
ES cells
ES derived cells
Heart
HSC & B–cell
IMR90
iPSC
Mesenchymal
Muscle
Myosatellite
Neurospheres
Other
Smooth muscle
Thymus
Chromatin state
Active
Inactive

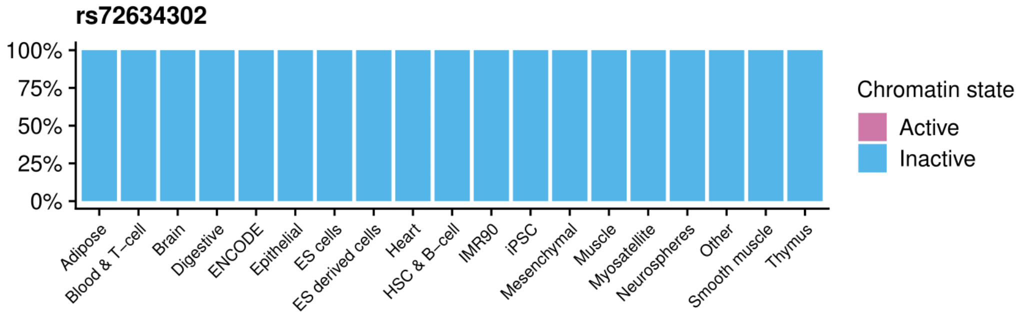
rs72634302
100%
75%
50%
25%
0%
Adipose
Blood & T–cell
Brain
Digestive
ENCODE
Epithelial
ES cells
ES derived cells
Heart
HSC & B–cell
IMR90
iPSC
Mesenchymal
Muscle
Myosatellite
Neurospheres
Other
Smooth muscle
Thymus
Chromatin state
Active
Inactive

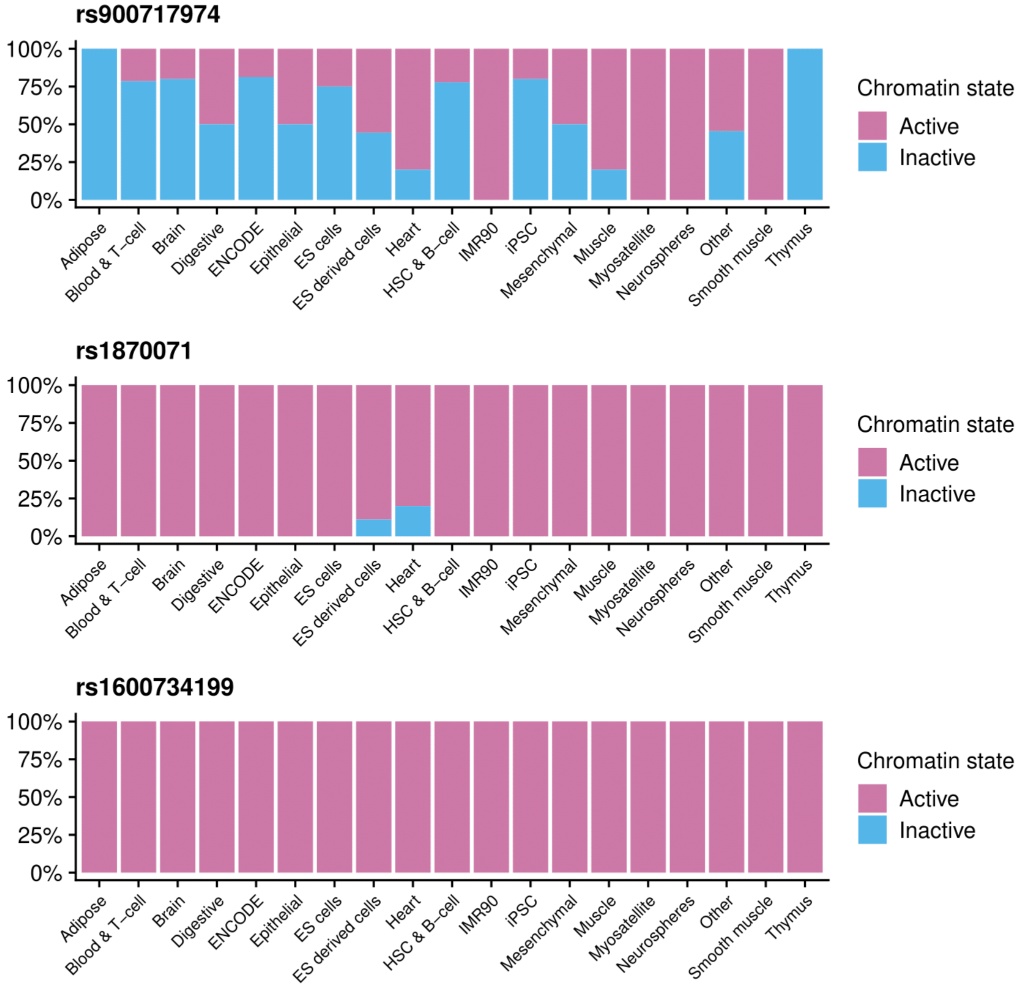

**Figure S6:** Chromatin state of the candidate SNPs of PNG lowlanders for the epigenomes of the Roadmap project clusters in tissues

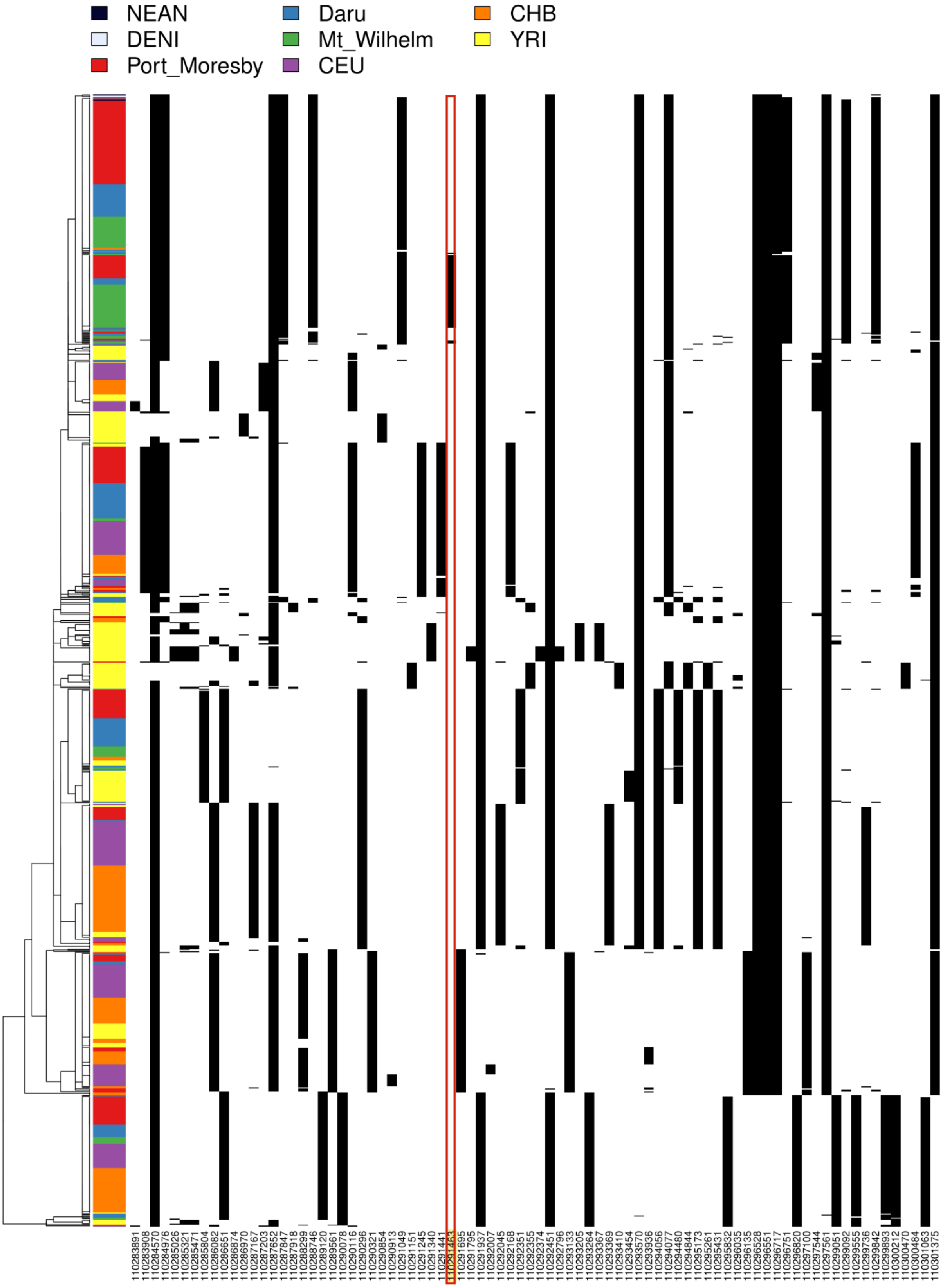

**Figure S7: Haplostrips plot centered on rs943845085, candidate SNP for the region chr4:110182324-110384099 under selection in PNG highlanders.** Introgression from Altai Neanderthal in PNG. The introgressed haplotype carry the SNP driving selection for the region (rs943845085, framed in red) but the Altai Neanderthal does not.

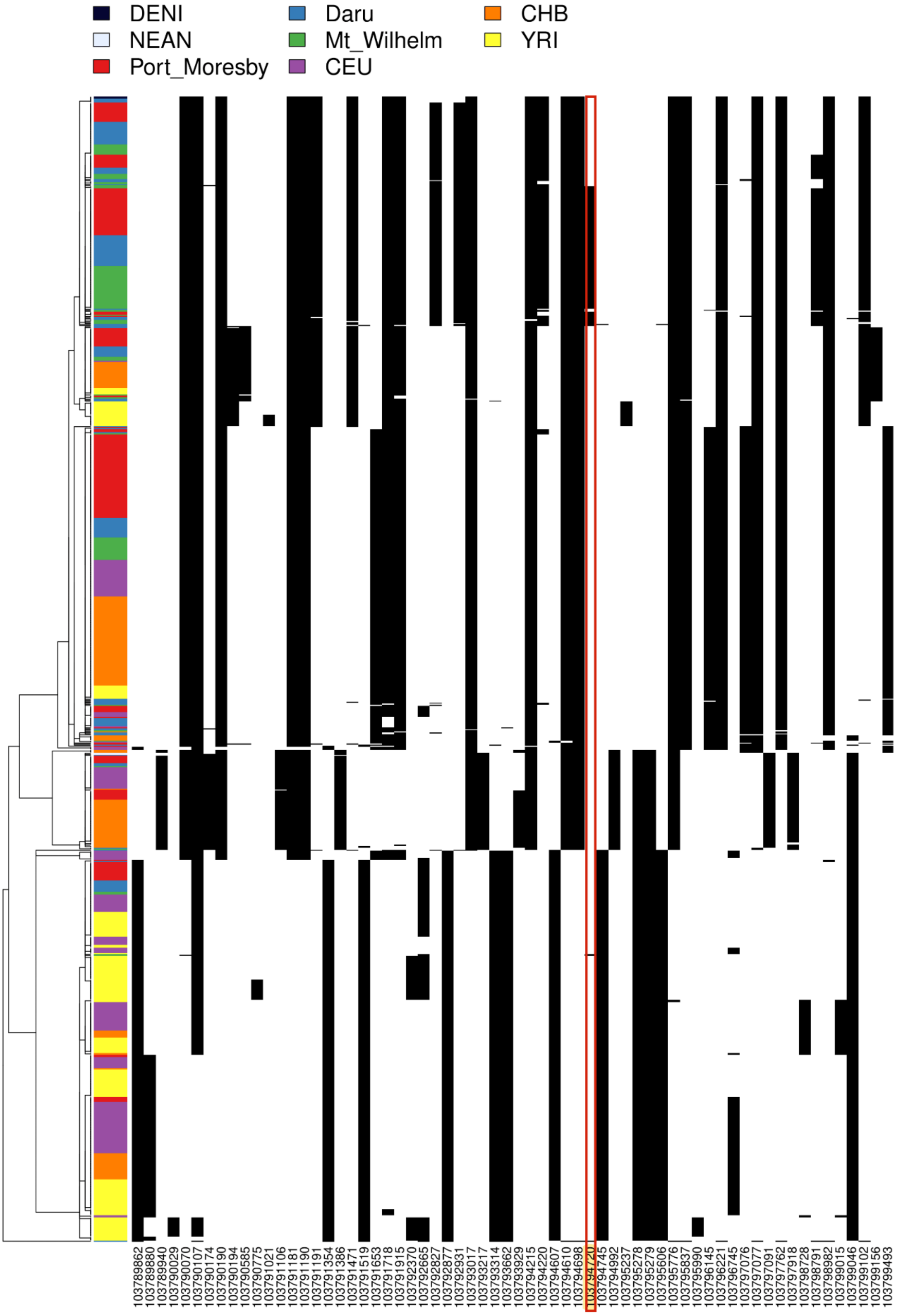

**Figure S8: Haplostrips plot centered on rs1032698711, candidate SNP for the region chr12:103783315-104121479 under selection in PNG highlanders.** Introgression from Altai Denisova in PNG. The introgressed haplotype carry the SNP driving selection for the region (rs1032698711, framed in red) but the Altai Denisova does not.

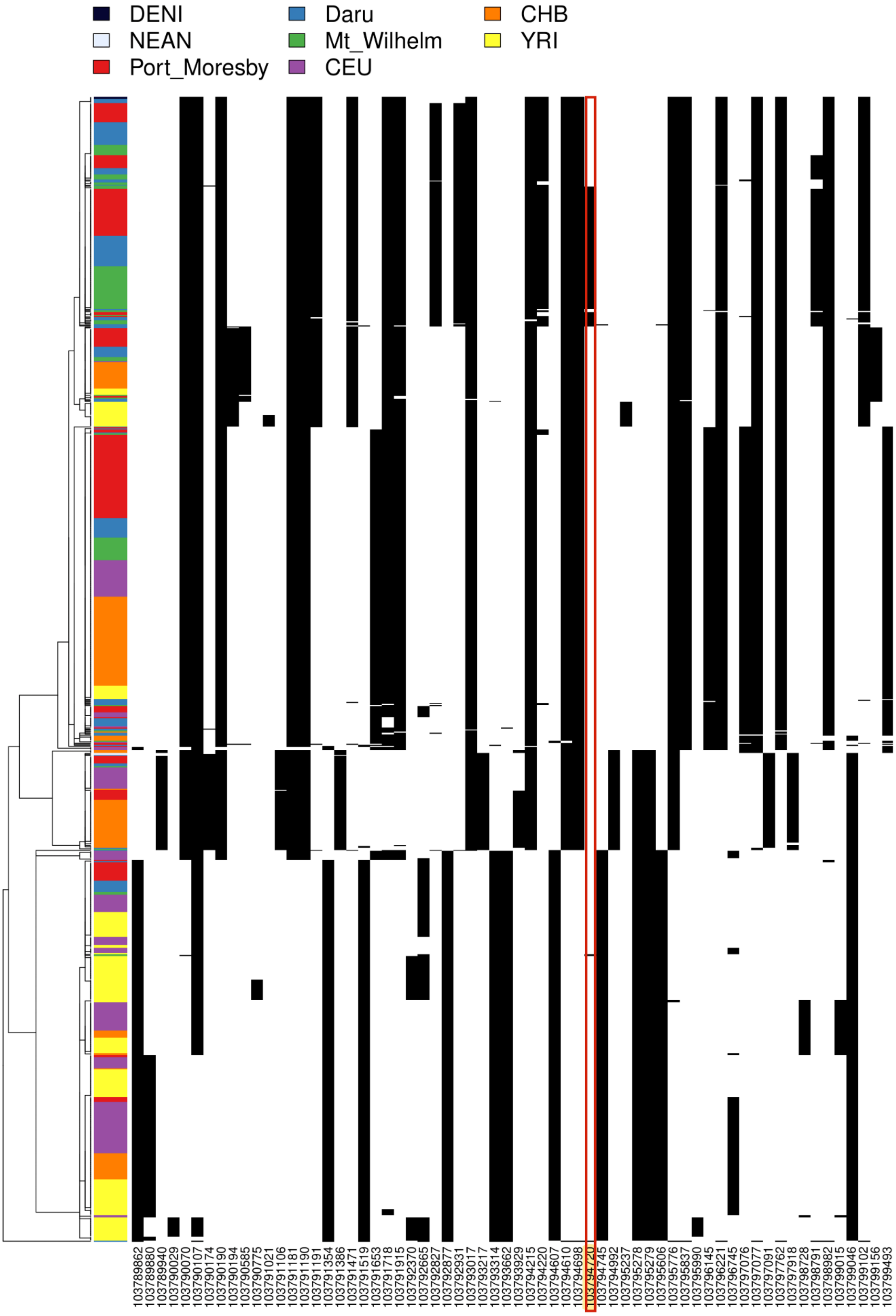

**Figure S9: Haplostrips plot centered on rs376150658, candidate SNP for the region chr2:241759136-242088831 under selection in PNG lowlanders.** Introgression from Altai Neanderthal in PNG. The introgressed haplotype carry the SNP driving selection for the region (rs376150658, framed in red) but the Altai Neanderthal does not.

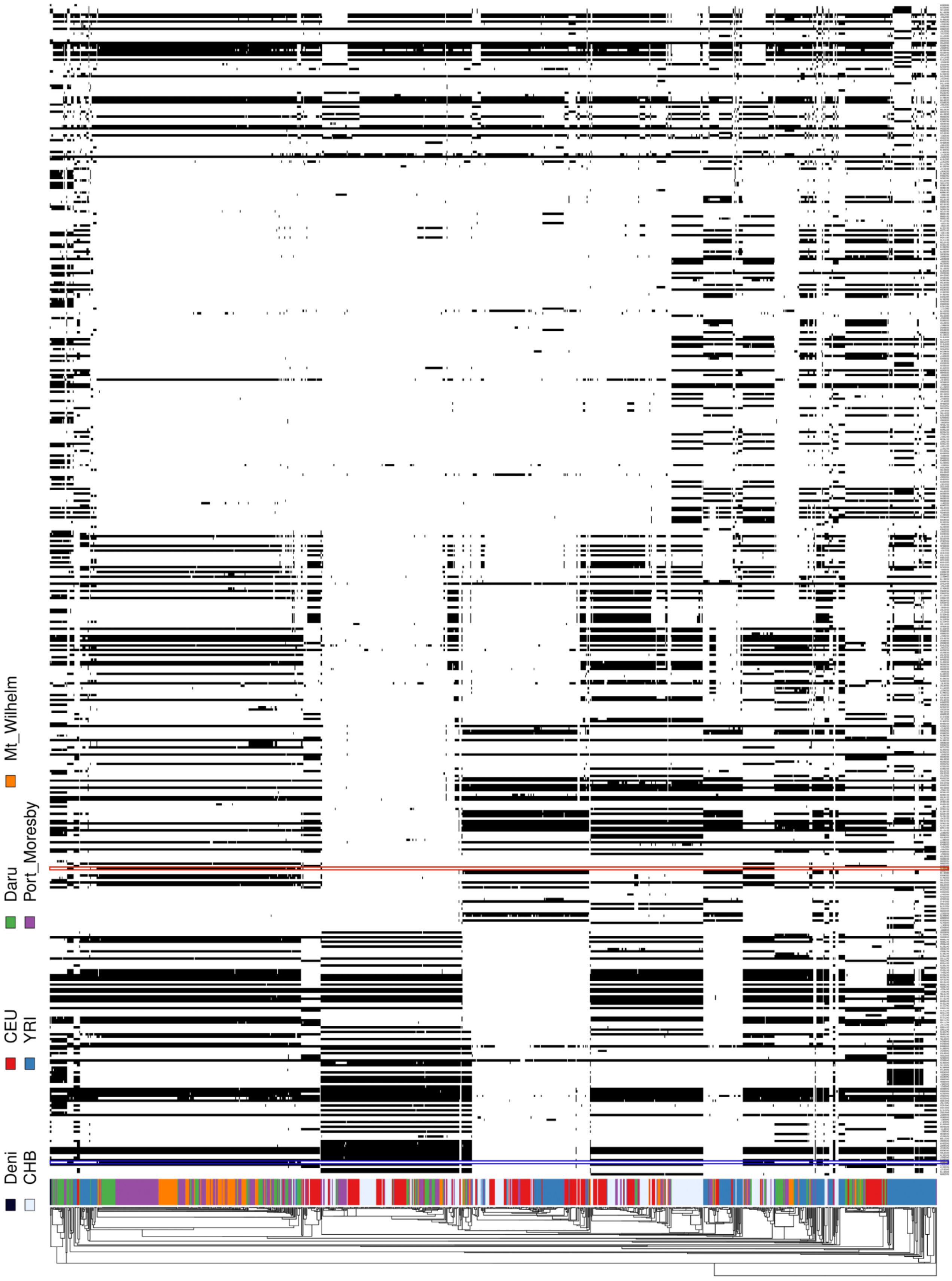

**Figure S10: Haplostrips plot for the region chr12:6452552-6662260 under selection in PNG highlanders.** Introgression from Altai Denisova in PNG. The introgressed haplotype does not carry the SNP driving selection for the region (rs74576183, framed in red) or the missense variant in LD with it (framed in blue).

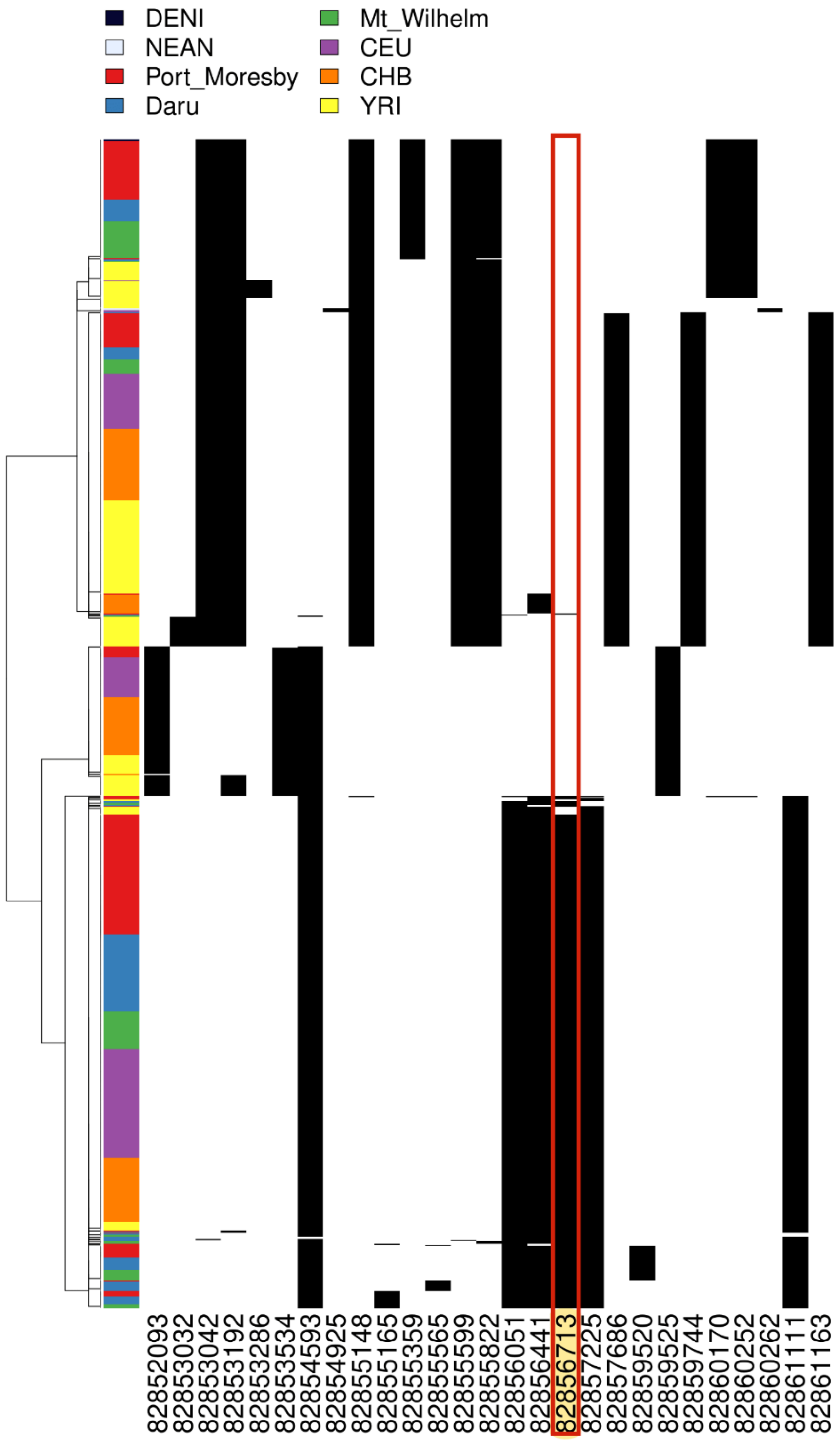

**Figure S11: Haplostrips plot centered on rs4693058, candidate SNP for the region chr4:82750503-83146792 under selection in PNG lowlanders.** Introgression from Altai Neanderthal in PNG. The introgressed haplotype carry the SNP driving selection for the region (rs4693058, framed in red) but the Altai Neanderthal does not.

### Supplementary Notes

#### 1. Sampling and sequencing

##### a. Sampling (with phenotypes)

For this paper, we sequenced 84 PNG individuals. These individuals were sequenced from saliva samples collected in two places in Papua New Guinea (PNG) between 2016 and 2019 (Table S1-S2). DNA was extracted from saliva samples with the Oragene sampling kit according to the manufacturer's instructions. Sequencing libraries were prepared using the TruSeq DNA PCR-Free HT kit. About 150-bp paired-end sequencing was performed on the Illumina HiSeq X5 sequencer. We also added 58 previously published PNG whole genome sequences [7]. These individuals were sampled in Port Moresby but are from various Papuan ancestry. Finally, we gained access to whole genome sequences from samples collected at the same sampling places during the same period and sequenced at the National Center of Human Genomics Research (France) (n=79) or KCCG Sequencing Laboratory (Garvan Institute of Medical Research, Australia)(n=41). These samples (n=262, PNG diversity II) are distributed between three sampling places, as show below:

- **PNG highlanders** (N=60): sampled in three different PNG highlands villages located on the slope of Mount Wilhelm (Chimbu Province): Womatne (2,300 m a.s.l.), Gembogl (2,500 m a.s.l.) and Keglsugl (2,700 m a.s.l.).
- **PNG lowlanders** (N=80): living below 100 m a.s.l. in Daru (Western Province).
- **PNG diversity I** dataset (N=122): individuals sampled in Port Moresby originating from various parts of PNG.

##### b. Kept Sequences

We detected six related pairs (Supplementary Note 6). When one individual of the related pair had a different number of phenotypes measurements, we kept the individual with the highest number of phenotype measurements. Or else we kept the individual with the highest mean coverage (Supplementary Note 4). We removed four mislabelled sequences from Mt Wilhelm. One sequence from Daru was removed before the variant calling (Supplementary Note 3). Finally, we removed two individuals with a low call rate (Supplementary Note 5).

##### c. Additional datasets (without phenotypes)

To these 262 PNG genomes, we added 81 published Papuan genomes (PNG diversity II) (Table S1) [2–6] and genomes from Yoruba in Ibadan, Nigera (YRI, n=108), Esan in Nigera (ESN, n=99) , British in England and Scotland (GBR, n=91), Utah Residents with Northern and Western European ancestry (n=99), Han Chinese in Bejing (CHB=103), China Kinh in Ho Chi Minh City, Vietnam (KHV, n=99) (The 1000 Genomes Project Consortium et al., 2015).

#### 2. Phenotype data collection

We chose the anthropometric and physiologic measurements based on reported adaptive mechanisms in highland populations (Beall 2013; Bigham and Lee, 2014). We explained the measurements to the study participants in detail. During the collection of the phenotypes, the participants were in the rested state, without shoes. We published these phenotypes measurements as a part of our dataset [10]. We had at least one phenotype measurement and age at the measurements for 234 of the 249 kept sequenced individuals (Table S3). We didn’t have phenotypes measurement for ten sequenced individuals (PNG lowlanders = 2, Port Moresby = 6, PNG highlanders = 2) and no age information for five individuals from Port Moresby. We didn’t include these individuals in the association analysis (Supplementary Note 17).

#### 3. Variant Calling

Sequencing data for all samples used in this study were processed together, starting from the raw reads. FASTQ files were trimmed with fastp v0.23.2 [11] using the default options, except for reads shorter than 35bp were discarded (-l 35), and auto-detection of adapters for paired-end sequence data was enabled (--detect_adapter_for_pe). Unpaired reads were removed. Filtered FASTQ files were converted to BAM using Picard Tools FastqToSam v2.26.2 [12]. Further processing was performed with Broad Institute’s GATK v4.2.0.0 Germline short variant discovery (SNPs and Indels) Best Practices [13]. HaplotypeCaller tool was used to produce individual sample GVCF files using Whole Genome Germline Single Sample v2.3.2 Cromwell workflow. The JointGenotyping workflow v1.5.1 was used to produce multi-sample VCF from GVCF files obtained in the first step. Data were processed with GRCh38 genome reference.

#### 4. Coverage

We computed mean coverage for the new sequences from PNG using the CollectWgsMetrics tool from Picardtools v2.26.3 within the autosomes interval (Table S4, Figure S1). The average mean coverage for all the PNG samples was 18.49x.

#### 5. Heterozygosity and call rate

We computed the heterozygosity and call rate of every sequence using plink (V 1.9) missing and het options. One PNG highlander (O517_DA_C001KNS) sequences and one PNG lowlander (191121_FD09254770) sequence were outliers for the call rate, with a call rate lower than 92%. We removed these two individuals from further analysis (Figure S2).

#### 6. Kinship analysis

We used KING v2.2.4 integrated relationship inference [14] to detect first and second-degree relatives pairs among our dataset using the option --related --degree 2. Within the 262 sequenced individuals, we detected six pairs of related individuals: 4 pairs of second-degree relatives (PNG highlanders = 1, PNG lowlanders = 3), one PNG lowlander parent-offspring pair and one full sibling pair from PNG diversity set I.

We kept the individual with the highest mean coverage from each pair (Supplementary Note 4). When two individuals from PNG diversity set I, Daru or Mt Wilhelm shared kinship but had a different number of phenotypes measurements, we kept the individuals with the highest number of phenotype measurements (Table S1).

#### 7. Filtering and Genomic mask generation

##### a. Generation of genomic mask

Previous publications used the 1000G genomic mask to indicate the sites with low accessibility to next-generation sequencing methods using short reads [15]. Because the 1000G genomic masks are based on low coverage, and our dataset is based on high coverage (both for high coverage 1000 genome data and our PNG data set), we computed a more adapted genomic mask.

Using cram files for the unrelated PNG highlanders, PNG lowlanders, PNG diversity set I individuals and 1000G populations of our dataset, we called the sites with a minimum base quality of 20, an alignment minimum mapping quality of 20 and downgrading mapping quality for reads containing excessive mismatches with a coefficient of 50. From these sites, we excluded indels, sites with more than two alleles, sites with a maximum missing rate of 5% and sites that weren't included in the mappability mask from MSMC (liftover to GR38) [16]. We also masked sites whose depth of coverage summed across all samples was higher or lower than the sum of the average depth across the dataset (24184.7) by a factor of 2-fold (below 12092.35 or above 48369.4). We wrote the remaining sites P, except for variants without the PASS flag from the variant calling.

##### b. Filtering

Unless otherwise stated, we performed the analysis stated in this paper on biallelic SNPs with a maximal missing rate of 5% filtered for our genomic mask. We excluded related individuals and two PNG samples with low call rate (Supplementary Notes 1,5,6 Table S1) from any further analysis.

#### 8. PCA

We performed Principal Component Analysis (PCA) using the smartpca program from the EIGENSOFT v.7.2.0 package [17] on LD pruned SNPs with a minor allele frequency (MAF) higher than 5%. To prune variants in high linkage disequilibrium, we used PLINK v.1.9 using the default parameters of 50 variants count window shifting from five variants and a variance inflation factor (VIF) threshold of 2 [18]. We used the R-3.3.0 software to plot the PCA. We computed the ten principal components (Figure S3).

#### 9. Admixture

We ran ADMIXTURE v1.3 [19] for the components K=2 to K=6 on the same dataset as the PCA (Supplementary Note 8). For each component, ADMIXTURE computes the cross-validation error using k-fold cross-validation procedure. We set the k parameter to 100. In order to select the model with the most likely number of components, we generated the cross-validation error 50 times for each component. We then defined the confidence interval of the cross-validation error for the component using the quantile of the 50 generated cross-validation errors. The lowest CV error is for K=5 (Figure S4). The confidence interval of the cross-validation error will be indicated as the following: (CI, 0.025 and 0.975 quantiles) for each component (Figure S4).

#### 10. Phasing

We phased sequences from Daru, Mt Wilhelm, PNG diversity set I and African, European and Asian populations from the 1000 Genome dataset. We used the software shapeit4 (v4.2.2) [20]. Because a reference appropriated to Papuans doesn't exist, we phased the samples statistically without reference. We used the genetic map provided by Eagle [21]. This phasing was performed per chromosome using the sequencing option that adjusts the default parameter to what the shapeit authors consider the most adapted to sequenced data.

#### 11. Population Branch Statistic (PBS)

We computed the Population Branch Statistic (PBS) distinctively on PNG highlanders and PNG lowlanders to detect recent natural selection signals. A population's PBS value represents the amount of allele frequency change at a given locus, and we are looking for the SNP that shows high differentiation on a branch of the population tree.

The allele frequency will be used at each locus to estimate the $F_{ST}$ between each pair of the three populations[22]. These pairwise $F_{ST}$ results will then be compared to estimate the change of frequency occurring in the target population since its divergence from the reference population [23].

We used respectively PNG lowlanders or PNG highlanders as the reference population and Yoruba population from 1000 Genome as an outgroup. We then defined sliding windows of 20 SNPs with a slide of 5 SNPs along the whole genome with bedtools (v2.29.2) [24]. We gave as PBS score for each of these SNP sliding windows the average PBS of the 20 SNPs inside the window. We then kept the windows whose average PBS score was in the 99$^{th}$ percentile and then merged the neighbouring left windows – that were not further than 10kb from each other - together. The PBS score of these merged windows is defined as the PBS score of the SNP windows with the highest average PBS score.

We considered the top 10 merged windows with the highest PBS score extended by a 50kb flanking region as the windows of interest for PBS (Tables S5, S6).

#### 12. Cross-extended haplotype homozygosity (XPEHH)

To explore a different kind of signature of positive selection in the two target populations, we used selscan (v2.0.0) to compute the cross-extended haplotype homozygosity (XP-EHH) on the phased dataset. XP-EHH is based on the reduced haplotype diversity that follows a selective sweep: selected allele and neighbouring allele frequencies will increase simultaneously due to linkage disequilibrium. [25].

We used PNG highlanders or PNG lowlanders as the target populations and, respectively, PNG lowlanders or PNG highlanders as the reference population and Eagle genetic map [21].

We kept the SNPs whose XPEHH score was in the 99$^{th}$ percentiles and then merged the SNPs not further than 10kb apart in a region together using bedtools. These regions were given as XPEHH score, the score of the SNP with the highest XPEHH score in the region.

We considered the ten windows with the highest XPEHH score extended by a 50kb flanking region as the windows of interest for XPEHH (Tables S5, S6).

#### 13. Fisher Score

We attributed to sliding windows of 20 SNPs (the same defined when computing PBS score) the XPEHH of the SNP with the highest XPEHH score of the 20 SNPs and the average of the PBS score of the 20 SNPs. We then combined these scores in a Fisher Score - the sum of the -log10 of the percentile rank for XPEHH and PBS [26] – for each sliding window. We kept only the windows with a Fisher Score in the 99$^{th}$ percentile that we then merged together when they were closer than 10kb with the closest windows. We kept the then merged windows, extended by a 50kb flanking region, with the highest Fisher Score as the regions of interest (Tables S5, S6).

#### 14. Relate

We computed recombination trees for the phased dataset using Relate (v1.1.8) [27].

We generated Relate input files from the phased dataset following Relate manual instructions and using a genomic mask we generated ourselves (Supplementary Note 8). With these inputs, we generated local trees using the default parameters for the mutation rate (1.25e-8 per generation per base pair) and the effective population size (30000) and Eagle genetic map[21]. We then extracted PNG highlanders subtrees and PNG lowlanders subtrees from the generated local trees.

Using the population-specific relative subtrees, we computed the coalescence rates through time separately for PNG highlanders and PNG lowlanders. We specified the first-time interval starting from 0 to $10^2$ years ago, and then the period between $10^2$ and $10^7$ is split into successive bins with a step of $10^{0.1}$. We used Relate default parameters of 28 years per generation, the number of cycles and the fractions of trees to be dropped.

Finally, we extracted the local trees for each SNPs in the regions of interest separately for PNG highlanders and PNG lowlanders and sampled branch length for those trees using an MCMC-like approach by applying the relevant Relate script. The branch length of the focal SNP tree is resampled 200 times, and each iteration result is recorded to estimate the uncertainty in the age estimate of each node.

To take the uncertainty in the branch length estimate into account, we performed this step 5 times. For each SNP of each region of interest, we thus computed sampled branches five times. We will run Clues for each of the five sampled branches.

#### 15. Clues

We used the CLUES [28], an approximate full-likelihood method for inferring selection, on each of the regions of interest previously characterized with XP-EHH, PBS and Fisher Scores (Supplementary Notes 10-12). This is to detect the SNP, which is the most likely to drive selection in each region.

CLUES uses the local trees generated for each SNP with Relate (Supplementary Note 13) in the regions of interest. The CLUES method will give each SNP the ratio of the likelihood of the SNPs being under selection – with a coefficient of selection computed by Clues – versus the likelihood of the SNP being under a neutral model. We computed the likelihood ratio for the five branch lengths generated for each SNP and averaged the five likelihood ratios. Finally, for each region, we selected the top five SNPs with the highest average likelihood ratios and generated branch lengths and logarithm ratios 50 additional times for these SNPs to further fine mapping for causal SNP (Tables S7-S10). Because SNPs with low DAF are unlikely to be under positive selection, we didn't consider SNPs with DAF lower than 5%.

#### 16. Association test

We used Genome-wide Efficient Mixed Model Association GEMMA (v0.98.4) [29] to detect if the SNPs likely to be under selection in PNG highlanders or PNG lowlanders are associated with any of the phenotypes we measured in the PNG population. We first created a centred relatedness matrix for all the unrelated PNG individuals for which we had at least one phenotype measurement (n=234) (Table S2). We will include this relatedness matrix in the univariate Linear Mixed Model (LMM) we used with GEMMA to correct the population structure. As we did in our previous study [10], we corrected height, diastolic pressure, systolic pressure, heart rate and haemoglobin for age and gender using a multiple linear regression approach. As height can be a covariable to some of these phenotypes, we also regressed chest width, waist circumference, FEV1, PEF and FVC, correcting for age, gender and height. For each phenotype, we performed the GEMMA LMM for the SNPs of interest and the phenotypes residuals with the relatedness matrix to control for population structure. Because we are performing the test for several SNPs, we corrected for multitesting with the Benjamini-

Hochberg procedure [30] for FDR < 5%. We corrected the p-value following the equation from [31]. For each phenotype, we multiply each p-value (one per SNP tested for the phenotype) per the total number of SNPs tested for the phenotype and divide it by the p-value rank order (smallest p-value for the phenotype = 1). We are also testing multiple phenotypes that gather in five groups of phenotypes highly correlated to each other [10] because the adjusted p-value should be considered significant when lower than 0.01 (0.05/5).

#### 17. Association in the UK biobank

To further understand how our candidate SNPs affect the phenotypes of the PNG populations, we downloaded the UK biobank's summary statistics[32] (Supplementary Note 18). We considered only phenotypes with a number of cases (n_cases) above 10,000 samples to work with common phenotypes. We are left with 1,931 phenotypes. Because our candidate SNPs are mapped on the Genome Reference Consortium Human Build 38 (GRCh38), and the UK biobank's summary statistics account for SNPs mapped on GRCh37, we converted our SNP list to GR37 using liftOver[33]. If the targeted SNP does not exist in the UK Biobank dataset, we chose the closest SNP from a 1kb upstream and 1kb downstream region. In order to avoid the ancestry sample size bias present in UKBB, we only extracted the p-value (pval_EUR) and beta score (beta_EUR) for European ancestry. After extracting the SNPs summary statistics for every phenotype, we removed all the phenotypes with -log(p-value) lower than -11.29 to correct for multitesting. We chose this threshold base on the standard significance p-value threshold in GWAS of $5 \times 10^{-8}$ [34] corrected for around the 1931 different phenotypes we used in the summary statistics (Tables S11-S15).

#### 18. Introgression

To reveal adaptive introgression between PNG haplotypes and archaic haplotypes, we used haplostrips (v1.3) [35] within PNG, African, Asian and European samples with Altai [36], Vindija [37], Chagyrskaya [38] Neanderthal or Denisovan [39] genome as reference haplotypes (Figure S5). We first subset Neanderthal and Denisova positions for the positions in our PNG and 1000 genome dataset with bcftools view (v.1.14) [40].

We ran haplostrips for the 21 regions under selection in PNG highlanders and the 23 regions under selection in PNG lowlanders using the phased dataset and the extracted Denisovan or Neanderthal positions. We used the haplostrips inner join option where only sites present in the PNG, the 1000G dataset and the archaic positions are used.
